# Phylogenomics supports a single origin of terrestriality in Isopods

**DOI:** 10.1101/2024.05.02.592215

**Authors:** Jessica A. Thomas Thorpe

## Abstract

Terrestriality, the adaptation to life on land, is one of the key evolutionary transitions, having occurred numerous times across the tree of life. Within Arthropoda, there have been several independent transitions in hexapods, myriapods, arachnids and isopods. Isopoda is a morphologically diverse order within Crustacea, with species adapted to almost every environment on Earth. The order is divided into 11 suborders with the most speciose, Oniscidea, including terrestrial isopods such as woodlice and sea-slaters. Recent molecular phylogenetic studies have challenged traditional isopod morphological taxonomy, suggesting that several well-accepted suborders, including Oniscidea, may be non-monophyletic. This implies that terrestriality may have evolved more than once within Isopoda. Current molecular hypotheses, however, are based on limited sequence data. Here, I collate available transcriptome and genome datasets for 33 isopods and four peracarid crustaceans from public sources, generate assemblies, and use 960 single-copy orthologues to estimate isopod relationships and the timing of divergences with molecular dating. The resulting phylogenetic analyses support monophyly of terrestrial isopods and suggest that conflicting relationships based on nuclear ribosomal sequences may be caused by long-branch attraction. Dating analyses suggest a Carboniferous-Permian origin of isopod terrestriality, much more recently than other terrestrial arthropods.

## INTRODUCTION

Isopoda is a large and morphologically diverse order within the arthropod class Crustacea, comprising over 10,000 species across 132 families and 11 suborders (Supplemental Figure 1) (1). The most speciose suborder is Oniscidea, the group containing woodlice and sea-slaters (>3,700 species; (2)). Adapted to almost every environment on Earth, isopods occupy a wide range of habitats; from tropical and temperate waters to polar regions. Marine isopods are found from the sublittoral zone to the deep-sea (3), and are associated with habitats such as seaweed forests (4), coral reefs (5), rocky shores, sandy sediments, and even hydrothermal vents (6). Freshwater isopods can be found in both open and subterranean water sources (7–9). Amphibious littoral isopods occupy shores and coastal regions, and while terrestrial isopods predominantly inhabit environments with high humidity, some are secondarily aquatic in both fresh and salt-water and others have adapted to life at high altitudes or even arid deserts (10,11). Isopods also vary substantially in size and feeding behaviour. The smallest isopods, at less than 0.2 mm, are meiofaunal (12), whereas giant, deep-sea isopods can reach up to 50 cm long (13). Terrestrial isopods are detritivores and browsers, feeding on decaying wood and vegetation. Many aquatic isopods fill a similar niche, feeding on dead and decaying plant and animal matter in the water. Some of these scavengers may also feed on slow moving sea creatures (14). Other marine isopods are filter feeders, sieving detrital and planktonic food particles from the water, while some are predators of smaller marine or fresh-water life. Several groups are parasitic, with different suborders adapted to different host phyla, such as fish or Crustacea (15–17).

Isopods are one of several groups of arthropods (alongside hexapods, arachnids, and myriapods) to have successfully colonised land. While the evolutionary route to terrestriality across arthropods is debated (*via* freshwater or littoral intermediates), the taxonomy of Oniscidea has long been considered a model of how terrestrial taxa may have evolved from marine ancestors. Woodlice possess many adaptations essential to non-aquatic life, for example, a water-resistant cuticle, carrying live young in a brood-pouch, a water-transport system and respiratory pleopods, as well as behavioural adaptations such as aggregation (18). Oniscidea has traditionally been divided into five groups, with the more derived terrestrial groups (Crinocheta, Synocheta, and Microcheta) possessing more complex adaptations that facilitate life on land. For example, within the Crinocheta, several families possess pleopodal lungs (two, three, or five pairs depending on the family) (19,20). In contrast, the ‘earliest’ terrestrial isopods (Tylida, Diplocheta) are predominantly littoral and display behavioural and morphological traits associated with aquatic life (21). For example, *Ligia*, a littoral sea-slater within Diplocheta, has a simple water-transport system, brood-pouch, and open respiratory pleopods, and has been hypothesised to represent the amphibious, marine ancestor of woodlice (22). Synocheta and Microcheta are considered intermediate, predominantly inhabiting highly humid environments, e.g. subterranean habitats (21,23).

The first phylogenetic analyses of Isopoda based on molecular data suggested that Oniscidea may not be monophyletic ((24,25,26); Supplemental Figure S2). In particular, these studies suggested that littoral oniscids may be more closely related to other marine suborders within Isopoda, and may have convergently adapted to life on land. For example, Lins et al. (25) collated publicly available isopod sequence data (nuclear ribosomal 18S, 28S, and mitochondrial cytochrome oxidase I (COI)) and recovered Oniscidea as paraphyletic. The two oniscid clades Diplocheta (comprising littoral genus *Ligia* and predominantly freshwater associated *Ligidium*) and Tylida (predominantly littoral) grouped with Phreatoicidea (Gondwana-distributed freshwater isopods), distinct from a clade containing the remaining Oniscidea (Crinocheta + Synocheta), albeit with extremely low phylogenetic support. Dimitriou et al., (26) investigated the relationships across the five oniscid superfamilies with 18S, 28S, and two nuclear protein-coding genes. Their analysis recovered *Ligia* alone outside Oniscidea, in a clade with other marine isopods, while the remaining members of Diplocheta and Tylida were placed in a monophyletic Oniscidea. The phylogeny of Dimitriou et al., (26) demonstrated robust phylogenetic support, however, sampling of non-terrestrial isopods was limited, so the closest marine relatives of *Ligia* (and potentially other members of Diplocheta and Tylida) as well as the relationships between other isopod suborders remain unresolved.

Across the other isopod suborders, molecular data has also indicated paraphyly of several accepted groups. Phylogenies based on nuclear ribosomal data have suggested non-monophyletic Asellota (the second-most speciose suborder including deep-sea isopods and freshwater pond-lice) (27), Cymothooidea (parasites of fish) (28), Sphaeromatidea (marine pill-bugs and sand-skaters) (29) and Anthuroidea (elongate, predatory ‘wormpods’) (25), in addition to Oniscidea. The position of the root among the earliest branching nodes within Isopoda is also unclear. Molecular dating analyses recovered one of the three parasitic clades, Cymothooidea, Epicaridea (crustacean parasites) or Gnathiidea (juvenile ‘pranzia’ are temporary parasites of marine fish, while sexually-dimorphic adults are non-feeding) near the root of the tree (25,28), whereas data from fossils and morphology place Asellota and Phreatoicidea as the earliest branching isopods, outside of ‘Scuticoxifera’; the clade comprising all the remaining suborders (1,30,31). Analyses of whole mitochondrial genomes have also been applied to isopod phylogenetics (32–34) recovering Asellota or Phreatoicidea as one of the earliest branching groups. However, factors such as nucleotide composition, GC-skew, and outgroup choice seem to impact which suborder is closest to the root in mitochondrial analyses, as well as affecting the monophyly of several other groups (Oniscidea, Epicaridea, and Cymothooidea) (32–34). Morphological studies have predominantly been in agreement regarding the monophyly of most suborders (with the exception of Oniscidea (35,36), and Cymothooidea (31,1)). However, relationships between suborders have been subject to multiple taxonomic revisions (see (1) for a summary of historical and recent studies). The most recent and perhaps well-known revision is the replacement of Flabellifera (comprising Cymothooidea, Sphaeromatoidea (marine pill-bugs), Seroloidea (sand-skaters) and Limnoriidea (tiny, marine wood-boring ‘gribbles’) with Cymothoida (consisting of Cymothooidea, Anthuroidea, Gnathiidea and Epicaridea, all demoted to superfamily level) (1). Furthermore, several morphologically disparate, taxon-poor suborders have not yet been placed confidently with either molecular or morphological data (e.g. Tainisopidea (7,8)). Indeed, some lack molecular data entirely (e.g. Phoratopidea, Calabozoidea (9,37)).

Genome and transcriptome data are now being deployed to resolve uncertain evolutionary relationships across the Tree of Life, including Arthropoda (38,39). While a comprehensive dataset does not yet exist for the whole of Isopoda, many isopod genomic and transcriptomic datasets are now publicly available. This study uses 960 single copy orthologues across 33 isopod genomic and transcriptomic datasets to infer a phylogeny for Isopoda. In contrast to previous molecular analyses, this phylogeny shows monophyly of Oniscidea, with a sister relationship between Diplocheta and Tylida, not observed previously. Other novel relationships recovered include the separation of the parasitic isopod clades Epicaridea and Cymothooidea. A separate analysis of traditional phylogenetic marker-genes indicates previous results may have been misled by long-branch attraction biases in nuclear ribosomal sequences. Molecular dating suggests that terrestrial isopods made the transition to land during the Carboniferous or Permian, much more recently than other terrestrial arthropods.

## METHODS

### Phylogenomic dataset

Genome and transcriptome datasets were collated for 33 isopod species, and for one amphipod, one cumacean and two tanaid outgroups. This dataset included 19 terrestrial oniscideans (13 Crinocheta, three Synocheta, two Tylida and one Diplocheta), and 13 marine and freshwater isopods (four Asellota, one Epicaridea, four Cymothoidea, two Valvifera, one Sphaeromatidea and one Limnoriidea) (Supplemental Table S1). Where data were available for more than one species in a given genus, the dataset with the highest gene completeness was selected (see below). FASTQ files were downloaded from the ENA database (www.ebi.ac.uk/ena).

Raw reads (Illumina RNA-seq and whole genome short-read sequences) were trimmed to remove low quality bases (quality cut-off score ‘-q 20’) and adapters (default parameters) with *fastp v0*.*23*.*2* (40). For genomic datasets, trimmed reads were assembled into contigs with *SPAdes v3*.*15*.*5* (kmer values, k = 21,33,55,77,99,127) (41). A scaffolded short and long read assembly for *Tylos granuliferus* was obtained from the authors (42). Transcriptome datasets were assembled into contigs with *Trinity v2*.*8*.*5* (43). The longest isoform for each gene was extracted with a custom python script (https://github.com/jessthomasthorpe/Isopod_Phylogenomics_MS). For each taxon, amino-acid and nucleotide sequences for single copy orthologs from the *arthropoda_obd10* set (comprising 1013 genes) were retrieved from the assembled contigs with *BUSCO v5*.*4*.*2* (Benchmarking Universal Single Copy Orthologs; (44)) (Supplemental Table S1). The python script *busco2fasta*.*py* (https://github.com/lstevens17/busco2fasta.py) was used to identify single-copy BUSCO genes and generate amino-acid and nucleotide fasta files for each. A cut-off of 50% taxonomic coverage was applied to each orthologue group, resulting in a total of 960 BUSCO sequences in the final dataset. To assess whether different alignment methods had any effect on isopod relationships, amino-acid BUSCOs were aligned with both *mafft v7*.*520* and *fsa v1*.*15*.*9* (45,46). Alignments were manually inspected to verify that detected isoforms were the same across taxa, else the least common isoform was removed from the dataset. To examine the effects of trimming, each set of alignments were either left untrimmed, or trimmed with *trimAl v1*.*4*.*rev15* (‘-gappyout’) (47). Nucleotide alignments were generated with ‘-backspan’ in *trimAl*, utilising the aligned amino-acid sequence as a guide (with and without trimming). This created a total of eight versions of the phylogenomic dataset for further analysis (Supplemental Data).

Phylogenetic relationships were examined across Isopoda using two different approaches to tree building. Firstly, a supermatrix approach; for each separate dataset, genes were concatenated with *catfasta2phyml*.*pl* (https://github.com/nylander/catfasta2phyml) and analysed by maximum likelihood (ML) in *IQ-TREE v2*.*2*.*0*.*3* (48,49). For the amino-acid datasets, an initial model selection was performed between four amino-acid substitution matrices (‘-m MFP -mset JTT, WAG, LG, Q.insect’), alongside various across-site variation and base frequency options available in *IQ-TREE* (50). This was followed by a preliminary analysis under the best-fit model (corresponding to the Q.insect matrix for each dataset, with estimated across-site rate variation parameters; Supplemental Table S2) to create a guide tree (51). A final thorough analysis with 1000 ultrafast bootstraps was run for each version of the dataset with the guide tree and selected rate variation parameters from the model-testing analysis, together with a C20 (20 component) profile mixture model in *IQ-TREE* (52,53). These posterior mean site frequency (PMSF) models in *IQ-TREE* are a rapid ML approximation to the CAT model of Le et al. (51), which allow for site specific variation in amino-acid preference across protein-coding sequences. For the nucleotide coding alignments, third codon positions were removed, then model selection followed by ML analysis under the best selected model was performed on each version of the dataset (Supplemental Table S2). To examine the effects of substitutional bias in synonymous sites, a ML analysis was also run for third codon positions, following the same procedure.

A second approach in phylogenetic analysis is to generate a summary species tree from individual gene trees, which incorporates any underlying discordance between individual genes. A summary species tree was calculated with *ASTRAL v*.*2*. Gene trees, and best model parameters for each gene, were estimated in *IQ-TREE2* (‘-m MFP -S’) (54). Congruence between the supermatrix tree and individual gene tree topologies was also assessed with concordance factors in *IQ-TREE* (55). Gene (GCF) and site (SCF) concordance factors (defined as the percentage of genes or sites that agree with the node in question) were calculated for each node from individual gene trees and sites in the alignment. Each node has four possible branching patterns: three different sister relationships, plus a polyphyletic arrangement, where the three taxa do not group together. GCFs were examined further to investigate the effect of removing the shortest (and potentially most noisy) genes from the analysis, on the proportion of polyphyletic gene trees for each node. The 960 genes were divided into ten sets, and GCFs recalculated ten times, removing the shortest 10% of genes each analysis. Finally, to establish whether the main topology was significantly preferred to any alternate topologies, ML hypothesis testing with the approximately unbiased (AU) test was performed in *IQ-TREE* (56). Alternative topologies included the *ASTRAL* summary tree, alternate placements of *Limnoria*, and topologies examining whether Diplocheta and Tylida were closer to marine isopods (Supplemental Figure S3).

### Marker-gene dataset

In addition to the phylogenomic dataset, a dataset of traditional phylogenetic marker genes was assembled to investigate the relationships observed in earlier isopod molecular studies. In total, 11 genes from published studies were collated across 148 isopods and 12 outgroups from Genbank (www.ncbi.nlm.nih.gov). These comprised nuclear ribosomal genes (18S and 28S), nuclear protein-coding genes (Na/K ATPase alpha subunit, NAK; phosphoenolpyruvate carboxykinase, PEPCK; histone subunit 3, H3; and intestinal fatty acid binding protein 2, IGFBP), mitochondrial ribosomal genes (12S and 16S), and mitochondrial protein-coding genes (COI, NADH dehydrogenase subunit 4, ND4, and cytochrome B, CYTB). In order to maximise coverage across this dataset, in some cases data from two species in the same genus (and in a few cases, the same family) were combined and treated as a single taxon. While the use of composite taxa is not optimal, it has precedence in super-matrix analyses, and simulations have shown it is better for phylogenetic estimation than a greater proportion of missing data (57,58). Care was taken to avoid generating composite taxa where genera might be non-monophyletic (Supplemental Table S3).

As base composition heterogeneity across taxa can add systematic error to analyses, particularly in third codon positions of protein-coding sequences (59,60), all codon positions were assessed for base heterogeneity (‘statefreq’) in *PAUP*4*.*0b10* (61). *PAUP\** uses a chi-squared test to compare observed versus expected base composition of sequences. Any partitions with significant heterogeneity were re-coded with RY-recoding (60); purines (G or A) were coded as R, and pyrimidines (C or T) as Y. Data were then retested; any RY-recoded codon partitions still suffering from significant composition bias were excluded (Supplemental Table S4). Chi-squared base composition tests were also calculated in *IQ-TREE* prior to phylogenetic analyses (48).

The publicly available sequences were supplemented by nuclear protein-coding sequences from the isopod genomes and transcriptomes (Supplemental Table S1). BLAST databases were built with *BLAST v2*.*13*.*0* ‘makeblastdb’ for contig sets, which were queried with isopod NAK, PEPCK, H3, and IGFBP sequences, and the top hit contig (e-value < 1e-50) selected for alignment. Coding sequences were extracted from longer genomic contigs with *MetaEuk v6*.*a5d39d9*, using translated marker-gene amino-acid sequences as reference (62). For H3, which is multi-copy, *Limnoria* had more than one significant top hit. While inspection of the alignment indicated third codon positions were not identical across these top hit sequences, RY-recoding rendered them identical, so the partition was retained.

Sequences were aligned in *MAFFT*, inspected for mis-alignments then manually adjusted. Hyper-variable regions in rRNA genes that could not be aligned across the dataset were excluded (Supplemental Data). Alignments were concatenated, and ML phylogenetic analyses were performed in *IQ-TREE* (48,49). An initial analysis was performed to determine the best substitution model for each partition of 18 codon positions and four ribosomal sequences (‘-q partition_file -m MFP’) (Supplemental Table S4) (63). This was followed by ML analysis under the best-fit model for each partition with 1000 ultrafast bootstrap replicates. For this dataset, Isopoda was constrained as monophyletic, due to the unexpected placement of several outgroup and one ingroup taxa. A separate ML analysis was also performed on the 18S alignment (under best fit model ‘SYM+R6’) to compare its topology against that of the whole dataset. Finally, to examine nucleotide composition and heterotachy (variable substitution rates between species) across sites in 18S, two further ML analyses were performed: one with RY-recoding, and another under the General Heterogeneous evolution On a Single Topology (GHOST) model of sequence evolution in *IQ-TREE* (64,65).

### Molecular dating

To obtain divergence estimates across the whole order, molecular dating was performed on the marker-gene dataset with taxonomic constraints from the phylogenomic dataset, in *BEAST v*.*1*.*10 (66)*. Prior distributions on the root and 15 other nodes were applied based on an interpretation of the isopod fossil record; full details of all prior distributions for fossil constraints and divergence times are given in Supplemental Table S5. Separate analyses with log-normal and uniform prior distributions were performed. To best estimate molecular dates for the five suborders without genomic data (i.e. those most subject to bias from 18S), phylogenetic inference in *BEAST* was also informed by topological constraints from the literature. For example, the placement of Phreatoicidea and Gnathiidea were informed by constraining ‘Scutocoxifera’ as monophyletic, which is supported by both morphology and mitochondrial genomes (30,31,34). A second assumed monophyly was that of Sphaeromatidea (Sphaeromatoidea and Seroloidea), also supported by morphological characters (1,30). To reduce the effects of nucleotide bias for Tainisopidea and Anthuroidea, which grouped together potentially erroneously in the ML tree, the preferred placement of each clade was estimated in separate unconstrained *BEAST* analyses, then a combined analysis run with preferred placements constrained (Supplemental Figure S4).

Estimated models of sequence evolution from the ML analysis were applied to each four-nucleotide coding partition (Supplemental Table S4). The HKY model of sequence evolution, with the transition/transversion ratio (kappa) fixed at 0.5, and gamma distributed rates across sites, was applied to each RY-coded partition. Analyses were run under a relaxed molecular clock, with the birth-death model of speciation, for 100 million generations (sampled every 10,000). Tree and clock parameters were linked across partitions (to avoid problems with over-parameterisation), and a separate mutation parameter, *Nu*, was estimated for each. The ucld.mean prior was set to a gamma distribution, with shape and scale parameters set to 0.001 and 1000, respectively. All other priors were default *BEAUTI v*.*1*.*10* values (67). Convergence and effective sampling were assessed in *Tracer v1*.*7* with all effective sample sizes >>100. A maximum clade credibility tree was constructed with *TreeAnnotator v*.*1*.*10* from 32,000 trees sampled in the posterior distribution from four converging runs.

## RESULTS AND DISCUSSION

### Phylogenomic dataset

Transcriptomes and genomes were assembled for 33 isopods and four other crustacean taxa as outgroups (Supplemental Table S1). These assemblies were surveyed for single-copy conserved loci (using the BUSCO arthropoda_odb10 set, (44)) present in at least 50% of taxa. The supermatrix alignment of amino-acid sequences from these 960 nuclear orthologues produced a fully resolved and well-supported phylogeny (Figure 1a, Supplemental Figure S5). Neither alignment method nor trimming had any effect on the topology, which was identical across all datasets with 100% bootstrap support at all nodes. Several suborders were recovered as monophyletic, including Asellota and Valvifera, which are both well-supported by morphology (68–70). The terrestrial isopods, Oniscidea (here comprising Tylida, Diplocheta, Synocheta and Crinocheta), were also recovered as monophyletic, in contrast to previous analyses based on nuclear ribosomal data (25,26); Supplemental Figure S1). Tylida and Diplocheta form a monophyletic group, which is sister to Crinocheta and Synocheta. These relationships differ from previous taxonomic studies of Oniscidea. The widely accepted phylogeny of Erhard (71) recovered Diplocheta as sister to the remaining four terrestrial groups, whereas Tabacaru & Danielopol (72) proposed Tylida as sister to the remaining clades, and Schmalfuss (73) recovered a sister relationship between Tylida and Crinocheta. Unfortunately, neither genomic nor transcriptomic data are yet available for Microcheta or Ligidiidae (Diplocheta), necessary for fully resolving evolutionary relationships within the suborder. Whether the sister relationship between Tylida and Diplocheta holds with the addition of more taxa (especially the addition of *Ligidium*) or more data (both members of Tylida have less than 50% of analysed orthologues) is not yet certain.

**Figure 1.**
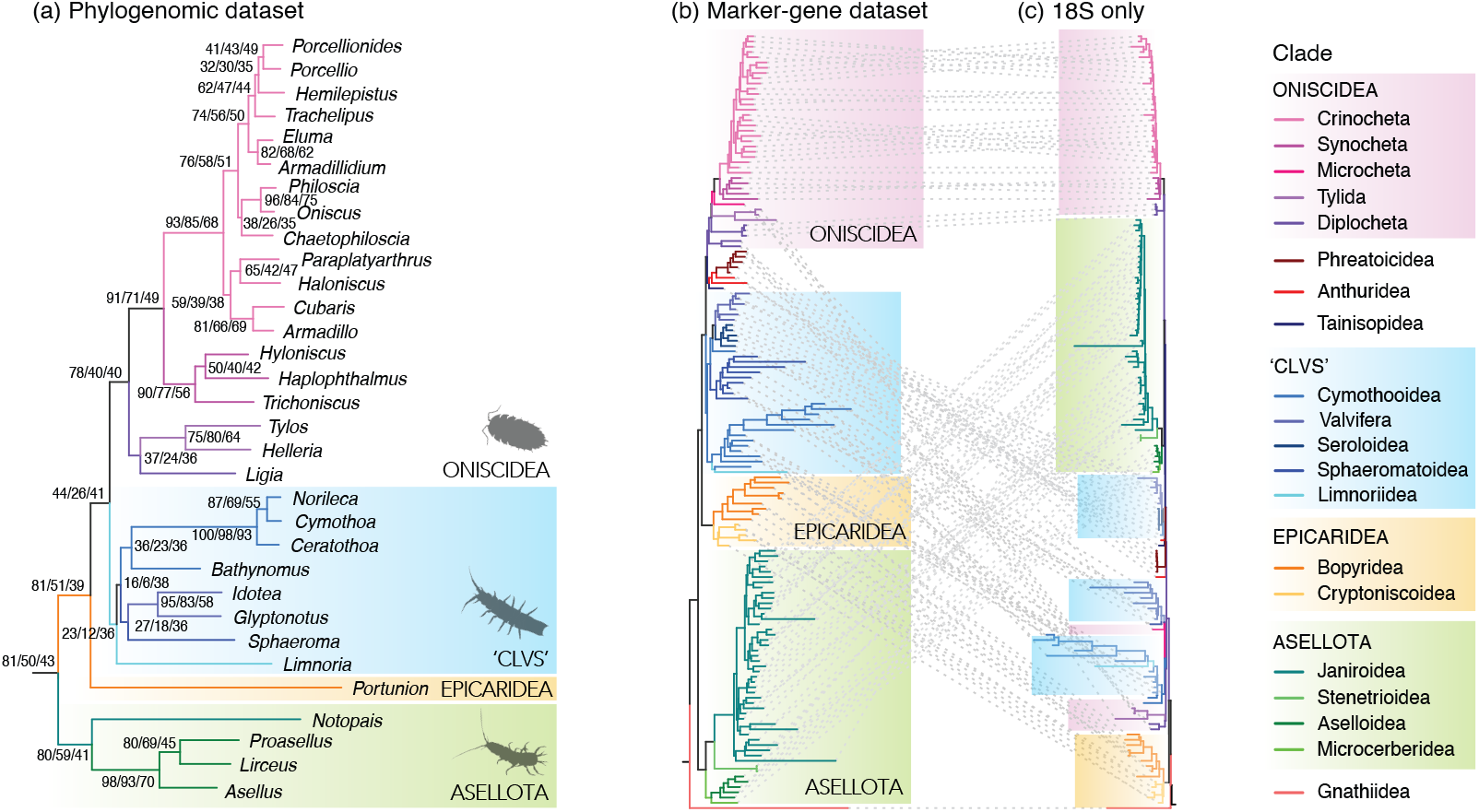
Phylogenies inferred by ML analysis of (a) the phylogenomic super-matrix of 960 single-copy orthologues and 33 species, aligned with *MAFFT* (b) the marker-gene super-matrix of 11 genes across 160 species and (c) 18S only from the marker-gene dataset. Nodal values in (a) indicate (R to L) concordance factors for sites (SCF), genes (GCF) and GCF for the longest 10% of genes (see main text). Full phylogenies are given in Supplementary Figures S5, S8 and S9.

Relationships within Crinocheta also differ from previous taxonomic studies (21). There are two well supported clades within Crinocheta; one consisting of predominantly northern hemisphere families (Porcellionidae, Agnaridae, Armadillididae, Trachelipodidae, and Oniscidae) together with the palaearctic philoscids, and the second comprising predominantly southern hemisphere families (Paraplatyarthridae, Armadillidae) and the antipodean philoscids. Taxonomists have long considered the family Philosciidae non-monophyletic, but the pattern in this study has not previously been reported (21). This well supported bifurcation is also observed in the marker-gene dataset, discussed below (Figure 1b).

Conversely, the suborder Cymothoida (comprising parasitic clades Cymothooidea, Epicaridea, Gnathiidea and predatory Anthuroidea; (1)) is not monophyletic; the two superfamilies present in this dataset, Cymothooidea and Epicaridea, are recovered separately in the tree (Figure 1a). This suggests there may have been at least two separate origins of parasitism in Isopoda, contrary to previous studies (1,30,74). Cymothooidea is monophyletic and is recovered in a clade with Limnoriidea, Valvifera and Sphaeromatoidea (hereafter referred to as ‘CLVS’), which together are sister to Oniscidea. Epicaridea is sister to this clade, and Asellota sister to all these, branching next to the root. These relationships also differ from previous molecular studies, where the two parasitic clades were recovered as sister groups (25,74). Instead, clade ‘CLVS’ is reminiscent of the original description of Flabellifera (68) (also referred to as ‘free-living flabelliferans’ in Brandt and Poore (1)), albeit with the inclusion of Valvifera. These results suggest the higher-level taxonomy of Isopoda may require revision; necessitating either the reinstatement of suborder status for Epicaridea (and Cymothooidea, Gnathiidea and Anthuroidea), or the creation of a new marine suborder containing Cymothooidea, Limnoridea, Valvifera and Sphaeromatidea (and potentially other groups) once more genomic datasets are available.

The sister relationship between Valvifera and Sphaeromatoidea in this analysis (no genomic data are yet available for Seroloidea) also differs from previous nuclear ribosomal phylogenies ((25,29) also Figure 1b), which recover Valvifera with Seroloidea, but Sphaeromatoidea with Cymothooidea, rendering Sphaeromatidea (Sphaeromatoidea and Seroloidea) non-monophyletic. In this phylogeny, Cymothooidea is sister to a clade comprising Valvifera and Sphaeromatoidea, and Limnoriidea is sister to these three groups. The relationships presented here thus lend support to the monophyly of Sphaeromatidea, a relatively recently proposed suborder, by Wägele (30) and recovered by the analysis of Brandt & Poore (1), though not Brusca & Wilson (31). The association between Limnoriidea and these marine suborders has also been observed in previous taxonomic studies, where Limnoriidea has been recovered with both Cymothooidea and Sphaeromatidea (1,30).

The topology of the species tree estimated from individual gene trees using *ASTRAL* is highly congruent with the supermatrix analysis (Figure 1a). The single difference is that Limnoriidea, rather than Cymothooidea, is recovered as sister to the Valvifera and Sphaeromatoidea clade (Supplemental Figure S6). The uncertainty in the placement of Limnoriidea between the *ASTRAL* and supermatrix phylogenies is reflected in the concordance factors. Gene (GCF) and site (SCF) concordance factors (given at nodes) indicate underlying topological variation, and are useful statistics for large genomic datasets, where bootstraps quickly reach 100% (55). Examining GCF and SCF for each node shows that while some relationships have little discordance (e.g. within Asellota), other nodes do not have such high underlying concordance (e.g. Oniscidea) and some are very discordant, with low GCF and SCF values (e.g. placement of Limnoriidea) (Supplemental Figure S7). Sequentially removing the shortest genes from the dataset (which potentially contain the least phylogenetic signal) increases GCF values and reduces the proportion of polyphyletic gene trees for almost all nodes across the phylogeny, including Oniscidea. While the proportion of polyphyletic gene trees is also reduced for the limnorid node, GCF remain low, suggesting there may be less signal for this node in the dataset. Tests of topology show a similar result (Supplemental Table S6); all alternate hypotheses except the topology produced in the *ASTRAL* phylogeny (where Limnoriidea is recovered as sister to the Sphaeromatoidea and Valvifera clade) were significantly rejected compared to the ML tree. Discordance or uncertainty at nodes might reflect real biological phenomena e.g. incomplete lineage sorting or ancient rapid radiation. If several isopod suborders diverged in quick succession over a short evolutionary time-frame, short internal branches with few informative substitutions may be overwritten by subsequent substitutions in long tips, obscuring phylogenetic signal and making phylogenetic resolution more difficult (75,76).

### Marker-gene dataset

ML analysis of a second dataset comprising 11 phylogenetic marker-genes (not present in the phylogenomic dataset), but with much denser taxonomic sampling, was also performed. This marker-gene dataset contains 148 isopods, across 9 of 11 suborders, with 12 outgroup taxa (Supplemental Table S2). The phylogeny from this dataset is remarkably similar to that of the phylogenomic dataset (Figures 1a and b, Supplemental Figures S5, S8 and S9) (and, interestingly, also similar to the most recent whole mitochondrial genome isopod phylogeny of Zou et al (34) (Supplemental Figure S1). In both phylogenies in this study, Asellota branches nearest the root, followed by Epicaridea, and both contain a clade comprising monophyletic Oniscidea and clade ‘CLVS’. However, there are also notable differences between the two datasets, specifically the relationships within clade ‘CLVS’. In the marker-gene dataset, *Bathynomus* (Cirolanidae) is not recovered with Cymothooidea but is instead sister to Seroloidea and Valvifera (with 100% bootstrap support), rendering both Cymothooidea and Sphaeromatidea non-monophyletic (Figure 1b).

To examine these relationships in more detail, a separate ML analysis was performed on only 18S, which is the longest gene in the marker-gene dataset, and present in all taxa (Fig 1c). Notably, it appears that areas of the marker-gene topology that differ to the phylogenomic dataset may directly reflect the 18S phylogeny, which appears to suffer from long-branch attraction artefacts. In the 18S phylogeny, clade ‘CLVS’ is split across the tree; *Bathynomus*, Serloidea and Valvifera form a strongly supported monophyletic clade nested within the phylogeny, whereas the remaining Cymothooidea, Sphaeromatoidea and Limnoriidea are recovered nearer the root, close to the parasitic groups Epicaridea and Gnathiidea, and the littoral oniscids Ligiidae (Diplocheta) and Tylida, as well as *Mesoniscus* (Microcheta). This topology therefore renders not only Cymothooidea and Sphaeromatidea non-monophyletic, but also Oniscidea; the remaining oniscids, Crinocheta, Synocheta and Ligiididae (Diplocheta) form a monophyletic clade sister to Asellota.

Robust testing for long-branch attraction, particularly in ribosomal sequences, is challenging (77). However, consistent with long-branch artefacts, is the observation that both the marine taxa and oniscids recovered near the root have much longer branch lengths than their closest relatives elsewhere in the tree (Figure 1; Supplemental Figures S8 and S9). In addition, many of the long-branched marine taxa near the root are parasitic, and parasites have been shown to have faster substitution rates (78), whereas the shorter-branched marine taxa nested in the tree are from deep-sea or cold-water habitats, which may result in slower rates (79). Finally, many relationships in the 18S phylogeny contradict not only the phylogenomic phylogeny but also well-accepted taxonomic relationships. This also suggests that the phylogenetic placement of several suborders lacking nuclear protein-coding sequences (and genomic or transcriptomic data) that are similar between the 18S and marker-gene phylogenies could be incorrect. These include a well-supported clade containing Anthuroidea, Tainisopidea and Phreatoicidea nested within the tree, and the placement of Gnathiidea at the root, or outside of Isopoda if the topology is unconstrained (1,7,30,31).

Incorrect phylogenetic estimation can arise due to sources of molecular bias (e.g., nucleotide composition or GC-skew in isopod mitogenomes; (33,34)). Chi-squared base composition tests indicate that while several members of Cymothooidea and one asellotan failed the individual composition test (Supplemental Table S3), there is no significant composition bias across the whole 18S dataset (e.g. unlike the third codon positions of protein-coding genes; Supplemental Table S4).

Furthermore, re-analyses with RY-recoding or under the GHOST model in *IQ-TREE* did not produce any meaningful changes to the topology (Supplemental Figures S10 and S11). However, ML analysis on third codon positions from the phylogenomic dataset (which can suffer from nucleotide biases such as saturation and base composition) produced a topology that was similar to the 18S dataset (Supplemental Figure S12).

These findings suggest that nuclear ribosomal sequences do not provide sufficient phylogenetic signal to evaluate higher-level isopod relationships. Furthermore, it may be the case that long-branch attraction in 18S sequences has been responsible for generating previous well-supported but ultimately incorrect higher-level phylogenetic groupings, including the recovery of non-monophyletic Oniscidea or a single origin of parasitism in isopods (25–29,74). While arthropod phylogenetics traditionally relied on nuclear ribosomal sequences prior to the widespread availability of genomic data, studies in insects have demonstrated that these sequences can be unreliable, the result of nucleotide composition biases and/or aberrant substitution rates (80,81). The underlying bias in isopod 18S sequences does not seem to result from differences in base composition (as it does in, e.g., isopod mitochondrial sequences), but rather from differences in substitution rate. Lifestyle and life history traits have been shown to affect species’ substitution rates e.g. (78,79,82,83) and multiple transitions between different environments and lifestyles over isopod evolutionary history could make differences in rates more pronounced.

### Molecular dating and biogeography of Isopoda

To examine the timing of various evolutionary transitions across Isopoda, a dated isopod phylogeny was estimated from the marker-gene dataset in *BEAST* with 16 calibrations from the fossil record (Figure 2, Supplemental Table S5). Topological constraints were applied from the phylogenomic dataset and taxonomic literature, meaning the isopod relationships estimated in this time-tree are slightly different to the marker-gene phylogeny (Figure 1b, Supplemental Figure S8). This approach was used to maximise the number of isopods in the dated analysis while limiting the effects of the 18S nucleotide bias. Estimated molecular dates were slightly older when prior distributions for fossil constraints approximated uniform, rather than lognormal distributions (Supplemental Figure S13). Placing isopod evolution in a temporal and biogeographic framework generates several interesting hypotheses regarding species distributions, vicariance and the timing of evolutionary transitions across the order (not just limited to terrestrial taxa).

**Figure 2.**
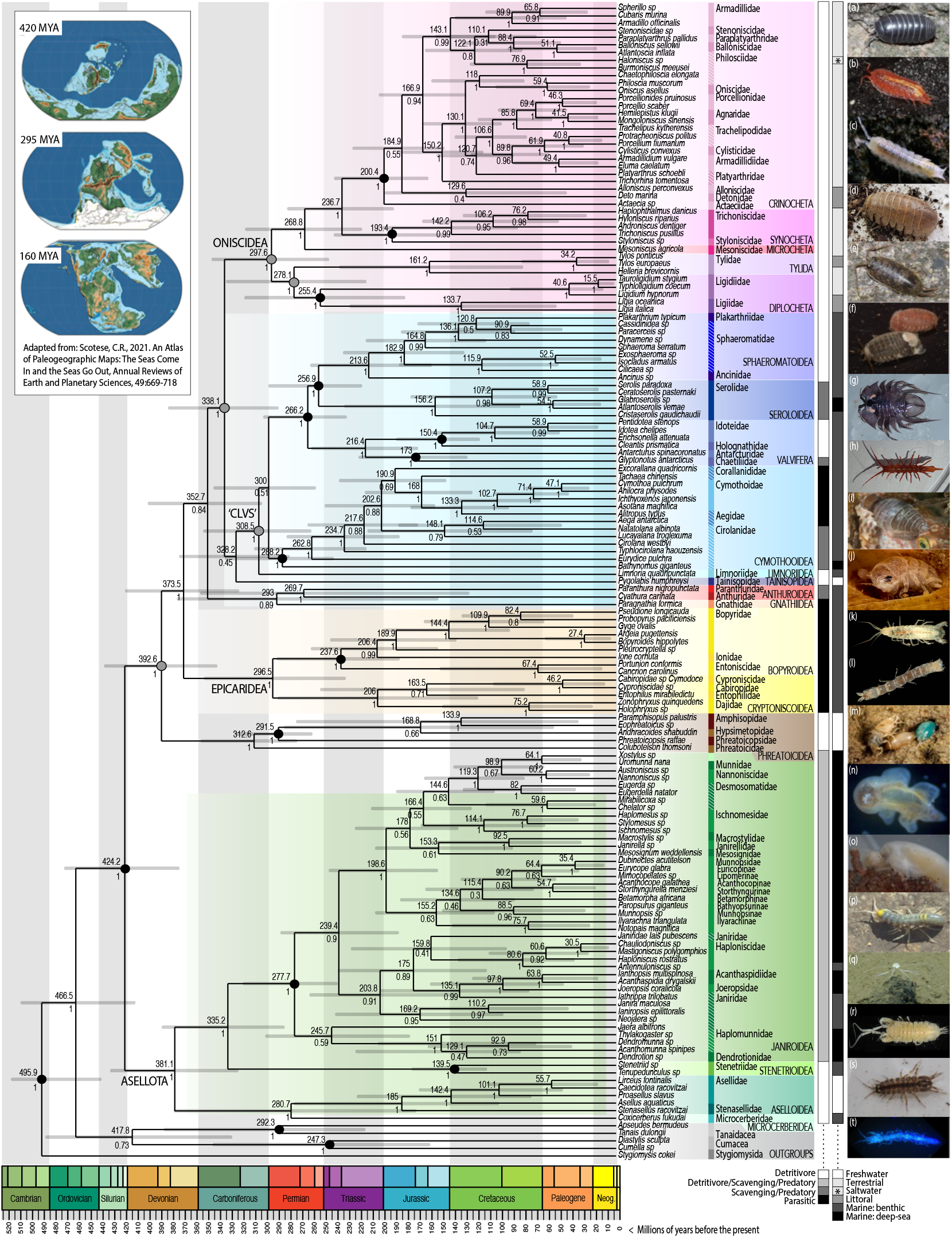
Dated phylogeny from *BEAST* analysis of 11 marker-genes with constraints from phylogenomic dataset (grey circles at nodes) and 16 fossils (black circles). Values at nodes indicate median divergence estimates in millions of years (above) and posterior probability (below). Current families are shown with vertical coloured bars, diagonal hatching indicates non-monophyly in this analysis. Predominant feeding behaviour and habitat are shown in greyscale bars. Inset left depicts three world maps at different time periods: late Silurian, 420 mya; early Permian, 295 mya; and late Jurassic, 160 mya. Images illustrate (a) Crinocheta, *Armadillidium vulgare* © W.Maguire; (b) Synocheta, *Androniscus dentiger* © W.Maguire; (c) Microcheta, *Mesoniscus graniger* © L.Kováč & P.Luptacik (d) Tylida, *Helleria brevicornis* © T.Hughes; (e) Diplocheta, *Ligia oceanica* © W.Maguire; (f) Sphaeromatoidea, *Sphaeroma serratum* © W.Maguire; (g) Seroloidea, *Brucerolis sp* © A.Hosie; (h) Valvifera, *Stenosoma lancifer* © J.Thomas Thorpe (i) Cymothooidea, *Anilocra sp* © S.Trewhella; (j) Limnoriidea, *Limnoria quadripuncata* © S.Trewhella; (k) Tainisopidea, *Tainisopus sp* © G.D.F.Wilson; (l) Anthuroidea, *Cyathura carinata* © S.Trewhella; (m) Gnathiidea, *Gnathia maxillaris* © S.Trewhella (n) Epicaridea, Cryptoniscoidea *Hemioniscus balani* © P.Adkins; (o) Epicaridea, Bopyroidea, *Athelges paguri* © J.Thomas Thorpe; (p) Phreatoicidea, *Phreatoicus sp*. © G.D.F.Wilson; (q) Janiroidea, *Munnopsidae sp*. © NOAA; (r) Stenetrioidea, *Stenetrium sp*. © G.D.F.Wilson; (s) Aselloidea, *Asellus aquaticus* © W.Maguire; (t) Microcerberidea, *Texicerberus sp*. © Benjamin Schwartz. (Further information in Supplementary Figure S1).

The earliest divergence in the dated phylogeny, between the mysids and mancoid peracarids, dates to ∼496 (446-521) million years ago (mya), between the late Ordovician and mid-Cambrian. The earliest divergence within Isopoda, between suborders Asellota and Phreatoicidea, dates to ∼424 (378-475) mya, between the Devonian and Ordovician. Divergence estimates across the tree suggest that while the earliest representatives of each suborder were likely around before the Permian mass extinction event ∼250 mya, most modern families diversified after this, as the Tethys Sea was newly forming (∼250 mya (84)) and Pangaea began to separate (∼200 mya; (85)).

### The earliest Isopoda: freshwater relicts and deep-sea radiations

Despite being one of the earliest-diverging isopod suborders, the asellote fossil record is sparse. The first asellote fossil constrains the appearance of the suborder in this analysis to at least the late Triassic (Supplemental Table S5), though Asellota are likely much older. Here, median divergence dates appear to be younger than previously published estimates for the suborder (25,28,86). The earliest divergence within Asellota, separating Aselloidea and Microcerberidea from Stenetrioidea and Janiroidea, dates to between the Silurian and Carboniferous, ∼335 (328-435) mya. The subsequent divergence between Microcerberidea and Aselloidea dates to ∼281 (174-380) mya, with biogeographic patterns aligning better to upper date estimates in the late Devonian or early Carboniferous. Microcerberidea are tiny interstitial marine and freshwater isopods with a Tethys-relict distribution. Marine species were likely once widespread along the shores of the Tethys Sea, before its regression during the Mesozoic, as demonstrated by the distribution of freshwater taxa in ground water sources once part of the ancient Tethyan shoreline (87,88). The divergence between Microcerberidea and Aselloidea should therefore be considerably older than the split of Laurasia and Gondwana (240-200 mya; (85)). It has been proposed that transitions between marine and freshwater habitats may have occurred many times in the evolution of Isopoda, through similar marine transgressions and regressions (early on, resulting in whole freshwater suborders e.g. Phreatoicidea, Tainisopidea, Calabozoidea but also more recently within certain families e.g. sphaeromatids or cirolanids (89)). In Asellota, transgressions may have occurred separately on both Laurasia and Gondwana; members of Aselloidea are almost all freshwater and found across the northern hemisphere, whereas freshwater Gnathostenetroidoidea have a southern hemisphere distribution (88). The common ancestor of all extant Aselloidea may be closer to upper divergence estimates, ∼185 (103-277) mya. However, within Asellidae, the divergence between North American (*Caecidotea* and *Lirceus)* and Eurasian (*Asellus* and *Proasellus)* species pre-dates the North Atlantic Split, ∼101 (47-169) mya.

Relationships within Janiroidea are largely consistent with previously published analyses, supporting four transitions to the deep-sea (28,90). The timing of deep-sea radiations in Isopoda has been debated, given that past oceanic anoxic events would have made the deep-sea uninhabitable at various points during the Mesozoic. It has been argued that deep-sea Asellota, however, may be evolutionarily ancient (91). The molecular dates estimated here suggest at least two of the transitions from shallow waters (in Acanthaspidiidae, ∼98 (17-123) mya and Haploniscidae, ∼81 (37-135) mya) may have occurred after the most recent anoxic events during the Cretaceous (‘Selli’ ∼120 mya and ‘Bonarelli’ ∼93 mya). Conversely, while the transitions to deep-sea life occurring in the ‘Munnopsid’ radiation (the largest radiation containing the majority of deep-sea families) ∼199 (155-247) mya, and the Dendrotionidae+Haplomunnidae radiation ∼151 (84-224) mya, likely occurred after the Permian mass extinction (an anoxic event ∼250 mya), both look to have radiated before anoxic events in the Cretaceous and potentially before the anoxic event in the Jurassic (Toarchian ∼183 mya), as previously proposed (91).

Unfortunately, this dataset cannot resolve whether Asellota or Phreatocidea are the earliest branching suborder of Isopoda. The earliest fossil isopods appear to be marine phreatoicids, with a wider distribution than today (92). The divergence estimates for Phreatoicidea may date the origin of these freshwater taxa from a marine transgression event in the southern hemisphere ∼313 (277-357) mya. Extant phreatoicids have a relict Gondwanan freshwater distribution, with species in New Zealand, Australia, India and South Africa likely dating to the break-up of Gondwana. For example, members of Hypsimetopidae can be found in both Australia and India, so the divergence within this family should date to at least 130 mya, when India separated from Gondwana (93). Estimates for the divergence between Amphisopidae and Hypsimetopidae, ∼169 (55-285) mya, appear to align with geological dates.

### Multiple origins of parasitism from free-living marine Isopoda

Divergence estimates for Scuticoxifera date between the mid-Carboniferous and Silurian-Devonian boundary, ∼374 (327-422). In this analysis, Epicaridea diverges before a clade containing Gnathiidea and Anthuroidea. In the traditional higher classification of Isopoda, Epicaridea, Gnathiidea and Anthuroidea were suborders, separate from Flabellifera (comprising Cymothooidea, Limnoridea and Sphaeromatidea) but recent taxonomic studies have all argued for a closer relationship between the parasitic suborders: Epicaridea, Gnathiidea, Cymothooidea, and Anthuroidea (1,30,31). Morphological characters have supported a sister relationship between both Gnathiidea and Anthuroidea (1), and Gnathiidea and Epicaridea (31). In this analysis, support for the sister relationship between Gnathiidea and Anthuroidea (0.89 posterior probability, PP) as well as placement of these groups outside of the Oniscidea+’CLVS’ clade (0.84 PP) is not particularly strong. Similar posterior probabilities are observed for the placement of Tainisopidea inside the Oniscidea+’CLVS’ clade in the unconstrained analyses (0.83 PP, Supplementary Figure S4), but equivocal for whether Tainisopidea is closer to clade ‘CLVS’ or Oniscidea. Previous taxonomic analysis has suggested that Tainisopidea might be related to Flabellifera, whereas the freshwater subterranean suborder Calabozoidea (currently without sequence data) may be closer to Oniscidea (88).

Fossil swellings, characteristic of infection by isopod parasites, can be seen in decapods from the Late Jurassic (Supplemental Table S5). In Epicaridea, the divergence between decapod parasites, Bopyroidea, and the parasites of other crustaceans, Cryptoniscoidea, dates to ∼296 (235-355) mya. The earliest crustacean parasites in Epicaridea may therefore have appeared around the same time as the earliest scavenging cirolanids in Cymothooidea, ∼288 (245-337) mya, but some time before the obligate fish parasites in Cymothoidae, ∼103 (67-141) mya. These latter dates suggest that cymothooid parasites may have evolved alongside their hosts, the teleost fish (between the early Triassic and late Cretaceous; (94)). In comparison, extant Epicaridea may have diverged sometime after the first appearance of decapods and other crustacean hosts in the Palaeozoic oceans (39,95).

The appearance of clade ‘CLVS’ dates to the Carboniferous, ∼309 (265-355) mya, and corresponds to the split between Limnoriidea and the remaining suborders (as in the phylogenomic dataset). Sphaeromatidea (Seroloidea and Sphaeromatoidea) and its divergence from Valvifera both date to the early Triassic. Valvifera and Sphaeromatoidea date to between the Permian and mid-Jurassic, ∼216 (172-262) and ∼214 (161-263) mya, respectively. That there are northern and southern hemisphere representatives in almost all families of both Sphaeromatoidea and Valvifera (except the valviferan arcturids which have been split into Arcturidae in the north, and Antarcturidae and related families in the south; (69)) suggest that both groups may have been widespread throughout the Tethys during the Triassic, and familial divergences should date to the breakup of Pangaea. These Triassic dates also contrast to divergence estimates, ∼156 (84-233) mya, (i.e. younger than the split of Pangaea) for Seroloidea, which has a Gondwanan distribution and likely origin (96–98). Within Serolidae, the divergence between South American and Antarctic serolids, ∼59 (16-116) mya, is slightly older than geological estimates for the opening of Drake’s Passage (∼49 mya (99)), which created a barrier for dispersal between the two continents.

### A single transition to land in terrestrial Isopoda

Dating the divergence of terrestrial isopods in Oniscidea also dates the origin of terrestriality in Isopoda. The earliest oniscid fossils date to the mid-Cretaceous, ∼105 mya (Supplementary Table S5). However, the earliest oniscid divergence in this analysis, between the littoral groups Diplocheta and Tylida and the remaining Oniscidea, dates to the Carboniferous-Permian boundary, ∼298 (249-348) mya. These dates suggest isopods made the transition to land considerably later than other terrestrial arthropods; molecular estimates for hexapods, myriapods and arachnids date to between the Ordovician (38) up to the Silurian or Cambrian (100), alongside the emergence of terrestrial plants.

Early oniscids may have been littoral, similar to *Ligia, Tylos* and the earliest branching families within Crinocheta (Scyphacidae, Actaeciidae, Alloniscidae, Detonidae). Evolutionary forays further inland therefore appear to have taken place multiple times, in each of the terrestrial groups. Outside of Crinocheta, terrestrial isopods are highly dependent on moisture, either through proximity to sources of surface water (e.g. riparian *Ligidium* (Ligidiidae) is associated with freshwater habitats), or through behavioural and physiological adaptations to life underground (e.g. *Helleria* (Tylida) lives in leaf-litter, but burrows below ground during droughts (101)). *Mesoniscus* and many Synocheta are subterranean, displaying adaptive, potentially convergent, traits for this lifestyle (21). The first transitions inland from littoral habitats may have occurred in the late Permian (potentially in northern Pangaea). *Mesoniscus* (Microcheta), found solely in Europe, is estimated to have diverged from Synocheta and Crinocheta ∼269 (221-319) mya. Within Diplocheta, the divergence between globally distributed Ligiidae and northern hemisphere Ligidiidae dates to ∼255 (184-319) mya. The divergence in Tylida between *Tylos* and forest-dwelling *Helleria* is more recent, ∼161 (82-238) mya.

In Synocheta, divergence estimates between southern hemisphere Styloniscidae and northern hemisphere Trichoniscidae ∼193 (148-242) mya, may date to the breakup of Pangaea. Within Crinocheta, the earliest branching littoral families are found across the southern hemisphere and tropics. The divergence between the predominantly northern and southern hemisphere clades dates to ∼167 (131-203) mya, which is younger than estimates in Synocheta. While this divergence could simply be underestimated, it is notable that no native northern hemisphere crinochetans are found in North America (102) (e.g. compared to the distributions of Asellidae, Ligidiidae, Trichoniscidae and many insect taxa) or throughout Asia (which has southern hemisphere Crinocheta), only more recent human-mediated introductions. This could suggest either localised extinction, or instead that Crinocheta may not have been widespread across Laurasia before it broke apart. Indeed, considering that many northern hemisphere crinochetan families are found in Europe and north Africa, these estimates might correspond better to the split of South America and Africa from the rest of Gondwana (which began ∼170 mya; (93)). However, the palaeobiogeography of Africa with Laurasia after separation from Gondwana is complex (103). Increased genomic and transcriptomic sampling across Crinocheta, particularly in Africa, may better elucidate the origins of northern hemisphere taxa, as well as other unexpected biogeographic patterns; e.g. the widespread Mediterranean distribution of *Armadillo officinalis*, a species whose closest relatives are found in the southern hemisphere. Increased sampling may also shed light on important aspects of oniscid evolution, such as the origin of pleopodal lungs. The phylogeny here suggests there could be a single origin in the northern hemisphere, ∼106 (78-138) mya, with the five more complex pleopods of Agnaridae and Trachelipodidae potentially evolving from five simple uncovered lungs, similar to *Oniscus* and *Philoscia* (19). Pleopods appear to have arisen several times in the southern hemisphere; in Armadillidae (potentially sharing an origin with Eubelidae, lacking sequence data), ∼86 (44-138) mya, and in southern hemisphere Philosciidae and Balloniscidae, ∼51 (18-92) mya (19,20). All these dates fall towards the latter part of the Cretaceous and early Palaeogene, which coincides with the climate becoming cooler and drier (104).

## CONCLUSIONS

Terrestrial isopods have long been considered an example of how marine taxa might transition to life on land. However, early molecular studies suggested multiple origins of terrestriality within Isopoda (25,26). In contrast to previous molecular phylogenies of Isopoda, this study finds compelling evidence for a monophyletic Oniscidea, as originally proposed by morphological analysis (21,30,31,71–73), and consistent with a single origin of terrestriality in isopods.

Importantly, this is the first study to use nuclear genomic and transcriptomic data to investigate and date evolutionary relationships across Isopoda, finding agreement between different phylogenetic approaches, different data types, and concordance across genes. Analysis of a taxonomically-rich marker-gene dataset indicates that previous studies may have been misled by long-branch attraction artefacts in nuclear ribosomal sequences. In addition, the relationships presented here are largely similar to those recovered in previous mitochondrial studies and a recent broader crustacean phylogenomic analysis (34,39). This study shows that coding sequences are both suitable and sufficient for resolving phylogenetic relationships in Isopoda, but greatly increasing taxonomic sampling with genomic data is essential to better understanding isopod evolution.

Isopoda have undergone numerous evolutionary transitions, enabling their radiation across almost all habitats on Earth. Sequencing across the order will enable a better understanding of not only the number and timing of these transitions; between marine and freshwater habitats, benthic and deep-sea ecosystems, free-living and parasitic lifestyles, and aquatic and terrestrial environments (and back again), but also which genes are involved. While there may have been just one transition to terrestrial life in Isopoda, pleopodal lungs have likely evolved several times. Genomic analysis will reveal whether similar transitions rely on the same underlying genetic machinery or if novel pathways are adopted. Through advances in sequencing technologies, and large-scale genomic projects, including the Darwin Tree of Life (105), high-quality genomes are increasingly becoming available. The generation of such genomes at scale for Isopoda will allow the evolutionary history and genetic basis of adaptation to diverse environments to be explored across the order.

