## Supplementary Tables for "Phylogenomics supports a single origin of terrestriality in Isopods"

**Supplementary Table S1**. Phylogenomic dataset, indicating taxonomic information; original sequencing information from NCBI Genbank, including sequencing method, SRA number and release date, instrument, reference (if published), as well as number of reads pre and post trimming with *fastp*; assembly information, including the number of scaffolds over 200 base pairs, with associated N50 value, longest scaffold length, assembly span and proportion of GC composition; and BUSCO information including the number and associated percentage of single copy, duplicate and fragmented BUSCO sequences from the *arthropoda_obd10* set.

| **Taxonomy** | | | | **Sequencing** | | | | | | | **Assembled Scaffolds > 200 bp** | | | | | **BUSCOs, n:1013** | | | | | |
|---|---|---|---|---|---|---|---|---|---|---|---|---|---|---|---|---|---|---|---|---|---|
| **Suborder** | **Superfamily** | **Family** | **Species** | **Method** | **SRA No.** | **Released** | **Instrument** | **Ref** | **# Reads** | **#Trimmed** | **#** | **N50** | **Max.** | **Span** | **GC** | **Sgl** | **%** | **Dupl** | **%** | **Frag** | **%** |
| Oniscidea | Crinocheta | Porcellionidae | *Porcellio scaber* | RNA-seq | SRR5298324 | 03/05/2017 | HiSeq 2000 | 1 | 8151981 | 7659870 | 26343 | 1119 | 55298 | 40.1Mb | 0.37 | 658 | 65.0 | 6 | 0.6 | 183 | 18.1 |
| Oniscidea | Crinocheta | Porcellionidae | *Porcellionides pruinosus* | RNA-seq | SRR15808558 | 08/09/2021 | HiSeq 2000 | 2 | 23941206 | 17388680 | 51643 | 956 | 16260 | 34.4Mb | 0.347 | 569 | 56.2 | 49 | 4.8 | 125 | 12.3 |
| Oniscidea | Crinocheta | Trachelipodidae | *Trachelipus rathkei* | RNA-seq | SRR24179284 | 06/06/2023 | HiSeq 2500 | | 60885658 | 58017691 | 119134 | 1346 | 24264 | 90.7Mb | 0.368 | 890 | 87.9 | 24 | 2.4 | 43 | 4.2 |
| Oniscidea | Crinocheta | Agnaridae | *Hemilepistus reaumurii* | RNA-seq | SRR26242025 | 30/09/2023 | NovaSeq 6000 | 3 | 92215308 | 92192876 | 646020 | 393 | 36634 | 269.5Mb | 0.413 | 935 | 92.3 | 16 | 1.6 | 35 | 3.5 |
| Oniscidea | Crinocheta | Armadillidiidae | *Armadillidium depressum* | RNA-seq | SRR5298308 | 03/05/2017 | HiSeq 2000 | 1 | 8970101 | 8663751 | 57300 | 1574 | 51269 | 50.4Mb | 0.364 | 884 | 87.3 | 11 | 1.1 | 72 | 7.1 |
| Oniscidea | Crinocheta | Armadillidiidae | *Eluma caelatum* | RNA-seq | SRR5298317 | 03/05/2017 | HiSeq 2000 | 1 | 7201366 | 6784539 | 70050 | 1198 | 44180 | 50.6Mb | 0.351 | 860 | 84.9 | 6 | 0.6 | 89 | 8.8 |
| Oniscidea | Crinocheta | Oniscidae | *Oniscus asellus* | RNA-seq | SRR5298319 | 03/05/2017 | HiSeq 2000 | 1 | 8444741 | 8201533 | 67534 | 1212 | 33538 | 51.3Mb | 0.367 | 804 | 79.4 | 10 | 1.0 | 118 | 11.6 |
| Oniscidea | Crinocheta | Philosciidae | *Philoscia muscorum* | RNA-seq | SRR5298320 | 03/05/2017 | HiSeq 2000 | 1 | 8381684 | 7906645 | 65129 | 1040 | 15534 | 47.0Mb | 0.357 | 826 | 81.5 | 11 | 1.1 | 109 | 10.8 |
| Oniscidea | Crinocheta | Philosciidae | *Chaetophiloscia elongata* | RNA-seq | SRR5298316 | 03/05/2017 | HiSeq 2000 | 1 | 8134572 | 7747047 | 68031 | 1259 | 26120 | 51.8Mb | 0.363 | 766 | 75.6 | 11 | 1.1 | 147 | 14.5 |
| Oniscidea | Crinocheta | Armadillidae | *Armadillo officinalis* | RNA-seq | SRR5298311 | 03/05/2017 | HiSeq 2000 | 1 | 7951577 | 7542693 | 57017 | 1007 | 23587 | 39.4Mb | 0.372 | 661 | 65.3 | 5 | 0.5 | 204 | 20.1 |
| Oniscidea | Crinocheta | Armadillidae | *Cubaris murina* | WGS | SRR14766320 | 09/06/2021 | NovaSeq 6000 | | 37454743 | 36352204 | 713651 | 1544 | 134018 | 724.3Mb | 0.339 | 168 | 16.6 | 12 | 1.2 | 390 | 38.5 |
| Oniscidea | Crinocheta | Philosciidae | *Haloniscus sp* | RNA-seq | SRR15808561 | 08/09/2021 | NovaSeq 6000 | 2 | 21354576 | 16916334 | 57534 | 1368 | 21586 | 48.9MB | 0.351 | 625 | 61.7 | 55 | 5.4 | 100 | 9.9 |
| Oniscidea | Crinocheta | Paraplatyarthridae | *Paraplatyarthrus sp* | RNA-seq | SRR15808560 | 08/09/2021 | NovaSeq 6000 | 2 | 24795047 | 18391224 | 65129 | 1040 | 15534 | 47.0Mb | 0.357 | 600 | 59.2 | 46 | 4.5 | 125 | 12.3 |
| Oniscidea | Synocheta | Trichoniscidae | *Haplophthalmus danicus* | WGS | SRR22764587 | 16/11/2023 | NovaSeq 6000 | 4 | 37833727 | 37723451 | 891792 | 809 | 840214 | 605.4Mb | 0.372 | 216 | 21.3 | 45 | 4.4 | 415 | 41.0 |
| Oniscidea | Synocheta | Trichoniscidae | *Hyloniscus riparius* | RNA-seq | SRR22938942 | 29/12/2023 | NovaSeq 6000 | 5 | 91669618 | 90708074 | 118165 | 1445 | 34076 | 93.4Mb | 0.378 | 893 | 88.2 | 28 | 2.8 | 45 | 4.4 |
| Oniscidea | Synocheta | Trichoniscidae | *Trichoniscus pusillus* | RNA-seq | SRR22938948 | 29/12/2023 | NovaSeq 6000 | 5 | 87804900 | 87135879 | 179467 | 1677 | 24718 | 152.8Mb | 0.379 | 934 | 92.2 | 31 | 3.1 | 24 | 2.4 |
| Oniscidea | Tylida | Tylidae | *Helleria brevicornis* | RNA-seq | SRR5298318 | 03/05/2017 | HiSeq 2000 | 1 | 8541441 | 7404707 | 34923 | 690 | 24110 | 19.6.Mb | 0.376 | 339 | 33.5 | 1 | 0.1 | 263 | 26.0 |
| Oniscidea | Tylida | Tylidae | *Tylos granuliferus* | WGS | DRR394944 DRR394945 | 13/12/2022 | HiSeq 4000 GridION | 6 | | | 200064 | 24713 | 615393 | 2.2Gb | 0.343 | 507 | 50.0 | 29 | 2.9 | 285 | 28.1 |
| Oniscidea | Diplocheta | Ligiidae | *Ligia exotica* | RNA-seq | SRR14289477 | 21/04/2021 | NovaSeq 6000 | 7 | 227871548 | 227116343 | 451889 | 844 | 36035 | 268.6Mb | 0.401 | 924 | 91.2 | 47 | 4.6 | 27 | 2.7 |
| Sphaeromatidea | Sphaeromatoidea | Sphaeromatidae | *Sphaeroma terebrans* | RNA-seq | SRR4436643 | 22/10/2016 | Analyzer IIx | 8 | 27757586 | 27358602 | 113037 | 1604 | 27683 | 99.7Mb | 0.446 | 857 | 84.6 | 64 | 6.3 | 55 | 5.4 |
| Valvifera | | Chaetiliidae | *Glyptonotus antarcticus* | RNA-seq | SRR15258043- SRR15258060 | 02/09/2021 | HiSeq 2500 | 9 | 674021026 | 664616893 | 283565 | 916 | 25035 | 179.3Mb | 0.402 | 862 | 85.1 | 55 | 5.4 | 46 | 4.5 |
| Valvifera | | Idoteidae | *Idotea baltica* | RNA-seq | SRR7056353 | 20/05/2020 | HiSeq 2500 | 10 | 296173922 | 279785070 | 113326 | 1500 | 27263 | 93.0Mb | 0.392 | 767 | 75.7 | 9 | 0.9 | 120 | 11.8 |
| Limnoriidea | | Limnoriidae | *Limnoria quadripunctata* | RNA-seq | SRR7059825 - SRR7059833 | 08/10/2018 | HiSeq 3000 | 11 | 377501758 | 373086579 | 2194578 | 326 | 25124 | 748.4Mb | 0.4 | 495 | 48.9 | 174 | 17.2 | 254 | 25.1 |
| Cymothoida | Cymothooidea | Cymothoidae | *Ceratothoa sp* | RNA-seq | SRR15808556 | 08/09/2021 | HiSeq 2000 | 2 | 24786465 | 24305480 | 44322 | 2305 | 35297 | 52.6Mb | 0.42 | 880 | 86.9 | 11 | 1.1 | 69 | 6.8 |
| Cymothoida | Cymothooidea | Cymothoidae | *Norileca indica* | RNA-seq | SRR18883214 | 01/03/2023 | NovaSeq 6000 | | | | 30894 | 2022 | 15279 | 38.0Mb | 0.411 | 933 | 92.1 | 14 | 1.4 | 21 | 2.1 |
| Cymothoida | Cymothooidea | Cymothoidae | *Cymothoa exigua* | RNA-seq | SRR8281006 | 07/12/2018 | NextSeq 500 | | 44238301 | 37978862 | 58944 | 1477 | 16702 | 48.0Mb | 0.436 | 896 | 88.5 | 10 | 1.0 | 58 | 5.7 |
| Cymothoida | Cymothooidea | Cirolanidae | *Bathynomus jamesi* | WGS | GCA023014485 | 15/04/2022 | PacBio Sequel | 12 | | | 22825 | 586500 | 4078738 | 5.9Gb | 0.373 | 775 | 76.5 | 158 | 15.6 | 43 | 4.2 |
| Cymothoida | Bopyroidea | Entoniscidae | *Portunion sinensis* | WGS | SRR15358545 | 11/08/2021 | BGISEQ-500 | | 28821067 | 28821061 | 495581 | 1130 | 57770 | 385.6Mb | 0.36 | 740 | 73.1 | 25 | 2.5 | 166 | 16.4 |
| Asellota | Janiroidea | Munnopsidae | *Notopais cryophila* | WGS | SRR16954879 | 31/12/2022 | NovaSeq 6000 | | 286555109 | 285238227 | 553909 | 4917 | 746767 | 1.0Gb | 0.305 | 480 | 47.4 | 19 | 1.9 | 322 | 31.8 |
| Asellota | Aselloidea | Asellidae | *Asellus hilgendorfii* | RNA-seq | SRR14135880 | 26/11/2021 | NextSeq 500 | 13 | 35917455 | 31902626 | 252227 | 1669 | 27733 | 209.4Mb | 0.373 | 944 | 93.2 | 21 | 2.1 | 15 | 1.5 |
| Asellota | Aselloidea | Asellidae | *Lirceus culveri* | RNA-seq | SRR11966488 | 01/01/2021 | HiSeq 2000 | 14 | 360958254 | 354411826 | 567774 | 1700 | 31936 | 434.2Mb | 0.365 | 882 | 87.1 | 70 | 6.9 | 29 | 2.9 |
| Asellota | Aselloidea | Asellidae | *Proasellus walteri* | RNA-seq | ERR3245438 | 25/03/2019 | HiSeq 2500 | 15 | 63282959 | 60782785 | 146897 | 2903 | 31818 | 174.3Mb | 0.346 | 927 | 91.5 | 17 | 1.7 | 18 | 1.8 |
| Tanaidacea | Apseudoidea | Apseudidae | *Apseudes sp* | RNA-seq | SRR14135879 | 26/11/2021 | NextSeq 500 | 13 | 29585462 | 24765781 | 110568 | 2368 | 28504 | 125.2Mb | 0.401 | 931 | 91.9 | 24 | 2.4 | 26 | 2.6 |
| Cumacea | | Nannastacidae | *Cumella sp* | RNA-seq | SRR5140154 | 01/02/2018 | HiSeq 2500 | 16 | 25181488 | 24732617 | 74550 | 605 | 13596 | 37.7Mb | 0.352 | 151 | 14.9 | 95 | 9.4 | 321 | 31.7 |
| Cumacea | | Diastylidae | *Diastylis cornuta* | RNA-seq | SRR25406671 | 24/07/2023 | HiSeq 2500 | 17 | 44120508 | 44020587 | 377293 | 835 | 48675 | 221.0Mb | 0.387 | 713 | 70.4 | 248 | 24.5 | 30 | 3.0 |
| Amphipoda | Gammaroidea | Gammaridae | *Gammarus electrus* | RNA-seq | SRR23685840 | 17/05/2023 | NovaSeq 6000 | 18 | 23167237 | 22922109 | 201996 | 1806 | 39535 | 181.0Mb | 0.46 | 703 | 69.4 | 239 | 23.6 | 40 | 3.9 |

**Supplementary Table S2.** Models selected with '-m MFP' in *IQ-TREE* for each version (alignment method and trimming) of the phylogenomic dataset; amino acid and nucleotide coding.

| Alignment | Trimmed | Length (amino acids) | Protein coding Model selected | Codon 1 ML model selected | Codon 2 ML model selected | Codon 3 ML model selected |
|---|---|---|---|---|---|---|
| ***MAFFT*** | **No** | 533909 | Q.insect, I+I+R6 | GTR+F+I+I+R8 | GTR+F+I+I+R7 | GTR+F+R8 |
| | **Yes** | 417813 | Q.insect, I+I+R6 | GTR+F+I+I+R9 | GTR+F+I+I+R6 | GTR+F+R8 |
| ***FSA*** | **No** | 675607 | Q.insect, F+I+I+R6 | GTR+F+I+I+R9 | GTR+F+I+I+R6 | GTR+F+R9 |
| | **Yes** | 403077 | Q.insect, I+I+R6 | GTR+F+I+I+R9 | GTR+F+I+I+R6 | GTR+F+R7 |

**Supplementary Table S3.** Marker Gene dataset indicating taxonomic information and accession numbers for genes included in this analysis from NCBI Genbank, except in a few cases for 16S and COI, which are from the BOLD database and indicated by '-XX', where XX is the release year. A * indicates a chimeric taxon, § indicates whether taxa failed individual 18S base composition test in *IQ-TREE* ML analysis. Accession AF255695, is in Genbank under *Cymodoce tattersalli,* however, based on its placement among Epicaridea in the phylogeny, it is likely a contaminant, maybe even an isopod parasite of Sphaeromatoidea.

| **SUBORDER** | | **FAMILY** | **GENUS** | **SPECIES** | **18S** | **28S** | **IGFBP** | **NAK** | **PEPCK** | **H3** | **16S** | **12S** | **CYTB** | **ND4** | **COI** |
|---|---|---|---|---|---|---|---|---|---|---|---|---|---|---|---|
| Oniscidea | Crinocheta | Porcellionidae | *Porcellio* | *scaber/dilatatus** | AJ287062 | EU914253 | MH880896 | Table S1 | Table S1 | Table S1 | DQ305104 | LC126630 | | KX289582 | LC126629 |
| Oniscidea | Crinocheta | Porcellionidae | *Porcellionides* | *pruinosus* | AY048181 | MN174809 | MH880898 | Table S1 | Table S1 | Table S1 | AJ275211 | KX467638 | KX289584 | KX289584 | KX289584 |
| Oniscidea | Crinocheta | Trachelipodidae | *Trachelipus* | *kytherensis/rathkei/ ratzeburgii ** | GQ302716 | MN174830 | Table S1 | MN234279 | MN234291 | Table S1 | EF027516 | KR013001 | KR013001 | MF187612 | KR013001 |
| Oniscidea | Crinocheta | Trachelipodidae | *Protracheoniscus* | *politus/fossuliger** | AY048184 | MN174817 | | MN234281 | MN234292 | | | | | | MG696252 |
| Oniscidea | Crinocheta | Trachelipodidae | *Porcellium* | *fiumanum* | AY048180 | | | | | | | | | | MT521289 |
| Oniscidea | Crinocheta | Agnaridae | *Hemilepistus* | *klugii* | MG887978 | MG888011 | | MG887926 | | | MG887951 | | | | MG887938 |
| Oniscidea | Crinocheta | Agnaridae | *Mongoloniscus* | *sinensis* | MW530521 | MW530520 | | MW534095 | MW534096 | | MG709492 | MG709492 | MG709492 | MG709492 | MW578682 |
| Oniscidea | Crinocheta | Cylisticidae | *Cylisticus* | *convexus* | AJ287059 | MN174813 | MH880886 | MN234280 | MN234293 | | KR013002 | KR013002 | KR013002 | KR013002 | KR013002 |
| Oniscidea | Crinocheta | Armadillidiidae | *Armadillidium* | *vulgare* | AJ267293 | AY739196 | MH880884 | | MN234299 | Table S1 | AF260847 | EF643519 | EF643519 | MF187614 | MH279710 |
| Oniscidea | Crinocheta | Armadillidiidae | *Eluma* | *caelatum/ purpurascens** | JN232925 | | MH880885 | | | Table S1 | AJ388100 | | | | MH279709 |
| Oniscidea | Crinocheta | Oniscidae | *Oniscus* | *asellus* | AF255699 | ON312047 | MH880887 | | | Table S1 | KX467635 | KX289581 | KX289581 | KX289581 | KX289581 |
| Oniscidea | Crinocheta | Philosciidae | *Philoscia* | *muscorum* | AJ287058 | | MH880889 | | | Table S1 | JF309310 | KX467636 | | | GU657075 |
| Oniscidea | Crinocheta | Philosciidae | *Chaetophiloscia* | *elongata* | MG887971 | | SRR5298316 | | | | MG887947 | | | | KJ668161 |
| Oniscidea | Crinocheta | Platyarthridae | *Platyarthrus* | *schoebli/ hoffmanseggi** | AJ287060 | MN174833 | MH880890 | MN234254 | MN234298 | | AJ388092 | KX467637 | | | KY020403 |
| Oniscidea | Crinocheta | Platyarthridae | *Trichorhina* | *tomentosa/officinalis** | AY048186 | | MH880905 | MN234282 | MN234300 | | JF309314 | OM326591 | | | MH295760 |
| Oniscidea | Crinocheta | Armadillidae | *Spherillo* | *dorsalis/grossus** | AB861916, KC999673 | LC496526 | | | | | KC706400 | AB861903 | | | KC706424 |
| Oniscidea | Crinocheta | Armadillidae | *Armadillo* | *officinalis* | GQ302704 | MN174812 | Table S1 | MN234252 | | Table S1 | AJ388094 | | OM304557 | | FN824106 |
| Oniscidea | Crinocheta | Armadillidae | *Cubaris* | *murina* | AJ287064 | | | | | | AB861890 | AB861895 | | | LC218701 |
| Oniscidea | Crinocheta | Philosciidae | *Haloniscus* | *sp* | KR424645 | | Table S1 | | | | | | | | EU364563 |
| Oniscidea | Crinocheta | Philosciidae | *Burmoniscus* | *meeusei/dasystylus** | AB889804 | AB889807 | | | | | AB889800 | AB626263 | | | AB889794 |
| Oniscidea | Crinocheta | Philosciidae | *Atlantoscia* | *inflata* | KY053446 | | MH880904 | | | | | | | | KJ814232 |
| Oniscidea | Crinocheta | Balloniscidae | *Balloniscus* | *sellowii* | KY706099 | | MH880901 | | | | | | | | KJ814235 |
| Oniscidea | Crinocheta | Alloniscidae | *Alloniscus* | *perconvexus/ oahuensis** | EU646199 | | | | | | ON980546 | ON980551 | | | OQ706274 |
| Oniscidea | Crinocheta | Stenoniscidae | *Stenoniscidae* | *sp* | KR424643 | | | | | | | | | | KR424605 |
| Oniscidea | Crinocheta | Paraplatyarthridae | *Paraplatyarthrus* | *pallidus* | KR424665 | | Table 1 | | | Table S1 | | | | | KR424578 |
| Oniscidea | Crinocheta | Scyphacidae | *Deto* | *marina* | KR424677 | | | | | | | | | | KR424585 |
| Oniscidea | Crinocheta | Actaeciidae | *Actaecia* | *sp/euchroa** | GQ302703 | MG888007 | | MG887930 | MN234324 | | GQ302691 | | | | GQ302701 |
| Oniscidea | Synocheta | Trichoniscidae | *Androniscus* | *dentiger/roseus** | JN232939 | MN174824 | | MN234283 | MN234313 | | | | | | BSOIL483-18 |
| Oniscidea | Synocheta | Trichoniscidae | *Haplophthalmus* | *danicus* | AJ287066 | | MH880908 | | | | KX467634 | KX467633 | | | MG311022 |
| Oniscidea | Synocheta | Trichoniscidae | *Hyloniscus* | *riparius/sp* | AJ287065 | | | | | | ON220031 | | | | MF743613 |
| Oniscidea | Synocheta | Trichoniscidae | *Trichoniscus* | *pusillus/provisorius** | AJ287067 | MN174835 | MH880909 | MN234259 | MN234316 | | AJ388088 | KX467642 | | | HQ966480 |
| Oniscidea | Synocheta | Styloniscidae | *Styloniscus* | *sp/magellanicus** | KR424679 | MN174832 | | | | | | KX467641 | | | KR424587 |
| Oniscidea | Microcheta | Mesoniscidae | *Mesoniscus* | *alpicola* | MN171513 | MN174829 | | MN234249 | MN234321 | | | | | | MT521129 |
| Oniscidea | Tylida | Tylidae | *Helleria* | *brevicornis* | GQ302709 | MN174843 | | MN234285 | MN234320 | Table S1 | GQ302690 | GU097465 | KJ468136 | | KJ468115 |
| Oniscidea | Tylida | Tylidae | *Tylos* | *ponticus/punctatus** | GQ302707 | MN174844 | | MN234265 | | KF007812 | KJ468185 | KJ468161 | KJ468144 | | EF027455 |
| Oniscidea | Tylida | Tylidae | *Tylos* | *europaeus* | EU646200 | | MH880910 | | | | KJ468169 | KJ468150 | KJ468131 | KF007343 | KJ468111 |
| Oniscidea | Diplocheta | Ligidiidae | *Tauroligidium* | *stygium* | MN171507 | MN174820 | | MN234255 | MN234305 | | | | | | ON212598 |
| Oniscidea | Diplocheta | Ligidiidae | *Typhloligidium* | *coecum* | MN171508 | MN174822 | | MN234251 | MN234309 | | | | | | ON212584 |
| Oniscidea | Diplocheta | Ligidiidae | *Ligidium* | *hypnorum/ghigii/lapetum** | AJ287056 | MN174818 | | MN234284 | MN234303 | JN800702 | JQ814403 | DQ182879 | | | DQ182781 |
| Oniscidea | Diplocheta | Ligiidae | *Ligia* | *italica* | GQ302705 | MN174838 | | MN234250 | MN234312 | | JQ814404 | DQ182954 | KF555753 | | KF555842 |
| Oniscidea | Diplocheta | Ligiidae | *Ligia* | *oceanica/exotica* | AY048178 | | MH880903 | | | MK034652 | JQ814402 | | DQ442914 | DQ442914 | KT208596 |
| Tainisopidea | | Tainisopidae | *Pygolabis* | *humphreysi* | GQ161216 | | | | | | | | | | EU364628 |
| Sphaeromatidea | Sphaeromatoidea | Ancinidae | *Ancinus* | *sp* | JF699514 | | | | | | KU248308 | | | | MH235844 |
| Sphaeromatidea | Sphaeromatoidea | Sphaeromatidae | *Isocladus* | *armatus* | JF699568 | | | | | | KU248302 | OK245257 | OK245257 | OK245257 | OK245257 |
| Sphaeromatidea | Sphaeromatoidea | Sphaeromatidae | *Sphaeroma* | *serratum/terebrans* | AF255694 | MN174841 | MH880907 | MN234262 | MN234301 | | KU248278 | GU130256 | MK460228 | MK460228 | GU130256 |
| Sphaeromatidea | Sphaeromatoidea | Sphaeromatidae | *Paracerceis* | *glynni* | AY743958 | AY739201 | | | | | KU248173 | | | | MH235892 |

| | | | | | | | | | | | | | | | |
|---|---|---|---|---|---|---|---|---|---|---|---|---|---|---|---|
| Sphaeromatidea | Sphaeromatoidea | Sphaeromatidae | *Cassidinidea* | *ovalis* | JF699522 | | | | | | KU248194 | | | | OQ322833 |
| Sphaeromatidea | Sphaeromatoidea | Sphaeromatidae | *Exosphaeroma* | *sp* | JF699543 | | | | | | KU248344 | | | | DQ889151 |
| Sphaeromatidea | Sphaeromatoidea | Sphaeromatidae | *Cilicaea* | *crassicaudata* | JF699524 | | | | | | KU248202 | | | | EF989646 |
| Sphaeromatidea | Sphaeromatoidea | Sphaeromatidae | *Dynamene* | *curalii* | JF699530 | | MH880906 | | | | KU248178 | | | | MK505687 |
| Sphaeromatidea | Seroloidea | Plakarthriidae | *Plakarthrium* | *typicum/punctatissimum** | JF699517 | | | | | | AJ269815 | | | | |
| Sphaeromatidea | Seroloidea | Serolidae | *Cristaserolis* | *gaudichaudii* | AJ269828 | | | | | | AJ269813 | | | | |
| Sphaeromatidea | Seroloidea | Serolidae | *Serolis* | *paradoxa* | AJ269827 | | | | | | EU419775 | | | | |
| Sphaeromatidea | Seroloidea | Serolidae | *Ceratoserolis* | *pasternaki* | AJ269826 | | | | | | AJ269801 | | | | EU597358 |
| Sphaeromatidea | Seroloidea | Serolidae | *Atlantoserolis* | *vemae* | KJ950677 | | | | | | KJ950662 | | | | KJ950683 |
| Sphaeromatidea | Seroloidea | Serolidae | *Glabroserolis* | *sp* | KJ950682 | | | | | | KJ950663 | | | | KJ950701 |
| Valvifera | | Arcturidae | *Antarcturus* | *spinacoronatus* | AF279604 | | | | | | AF268206 | | | | AWIAM135-09 |
| Valvifera | | Holognathidae | *Cleantis* | *prismatica* | AF255697 | JQ425584 | | | | | | | | JQ425553 | JQ425511 |
| Valvifera | | Chaetiliidae | *Glyptonotus* | *antarcticus* | AF255696 | | Table S1 | Table S1 | Table S1 | Table S1 | AM086486 | GU130254 | GU130254 | GU130254 | GU130254 |
| Valvifera | | Idoteidae | *Idotea* | *chelipes/wosnesenskii/ baltica** | GQ302710 | KC428847 | Table S1 | MN234264 | MN234310 | Table S1 | GQ302689 | AF260560 | DQ442915 | DQ442915 | DQ442915 |
| Valvifera | | Idoteidae | *Erichsonella* | *attenuata* | AY743948 | AY739191 | | | | | | | | | MH087590 |
| Valvifera | | Idoteidae | *Pentidotea* | *stenops* | JN800581, JN800618 | | | | | | JN800552 | JN800524 | | | MH242909 |
| Limnoriidea | | Limnoriidae | *Limnoria* | *quadripunctata* | AF279599 | FJ611321 | Table S1 | Table S1 | Table S1 | Table S1 | KF704000 | KF704000 | | KF704000 | KF704000 |
| Cymothoida | Anthuridea | Anthuridae | *Cyathura* | *carinata/sp** | AF332146 | KC428835 | | | | | AJ388072 | | | | AF520440 |
| Cymothoida | Anthuridea | Paranthuridae | *Paranthura* | *nigropunctata /japonica** | AF279598 | | | | | | GQ302694 | | | | ZPC614-18 |
| Cymothoida | Cymothooidea | Aegidae | *Aega* | *antarctica/sp** | AF255689 | | | | | | LC159470 | | | | FJ581463 |
| Cymothoida | Cymothooidea | Aegidae | *Alitropus* | *typus* | KT823938§ | | | | | | | | | | KT445864 |
| Cymothoida | Cymothooidea | Cymothoidae | *Anilocra* | *physodes* | AF255686§ | | | | | | EF455809 | | | | MK652476 |
| Cymothoida | Cymothooidea | Cymothoidae | *Asotana* | *magnifica* | MK542856§ | | | | | | MK790137 | MK790137 | | MK790137 | MK790137 |
| Cymothoida | Cymothooidea | Cymothoidae | *Ichthyoxenos* | *japonensis* | MK542857§ | | | | | | KX671993 | MF419233 | | MF419233 | MF419233 |
| Cymothoida | Cymothooidea | Cymothoidae | *Cymothoa* | *pulchrum/indica** | MH395141§ | | Table S1 | Table S1 | Table S1 | Table S1 | LC159450 | MH396438 | | MH396438 | LC159561 |
| Cymothoida | Cymothooidea | Corallanidae | *Tachaea* | *chinensis* | MK542858 | | | | | | LC160308 | MF419232 | | MK007965 | MF419232 |
| Cymothoida | Cymothooidea | Corallanidae | *Excorallana* | *quadricornis* | AF255688 | | | | | | | | | | |
| Cymothoida | Cymothooidea | Cirolanidae | *Eurydice* | *pulchra/affinis** | AF255690 | | | | | | AJ388073 | GU130253 | CO157399 | GU130253 | GU130253 |
| Cymothoida | Cymothooidea | Cirolanidae | *Cirolana* | *westbyi/rugicauda** | MK834589 | | | | | | AF259544 | AF260558 | | | AF260840 |
| Cymothoida | Cymothooidea | Cirolanidae | *Lucayalana* | *troglexuma* | KY426830 | | | | | | KY426828 | | | | KY426818 |
| Cymothoida | Cymothooidea | Cirolanidae | *Natatolana* | *albinota/rossi** | AF255691 | | | | | | GQ302693 | | | | MH509756 |
| Cymothoida | Cymothooidea | Cirolanidae | *Typhlocirolana* | *haouzensis* | AF453249 | | | | | | AF356848 | FJ460466 | | | |
| Cymothoida | Cymothooidea | Cirolanidae | *Bathynomus* | *giganteus/jamesi** | SRR8705862 | | JAJOZX01 | JAJOZX01 | JAJOZX01 | JAJOZX01 | MG229279 | KU057374 | KU057374 | KU057374 | MN654917 |
| Cymothoida | Cymothooidea | Gnathiidae | *Paragnathia/ Gnathia** | *formica* | AF255687 | | | | | | | | | | MT186550 |
| Cymothoida | Epicaridea: Bopyroidea | Bopyridae | *Probopyrus* | *pacificiensis* | AF255683 | | | | | | | | | | MK308333 |
| Cymothoida | Epicaridea: Bopyroidea | Bopyridae | *Pseudione* | *longicauda* | KF765760 | | | | | | | | | | LC476591 |
| Cymothoida | Epicaridea: Bopyroidea | Bopyridae | *Bopyroides* | *hippolytes* | MW540884 | | | | | | MW540884 | | | | MK905237 |
| Cymothoida | Epicaridea: Bopyroidea | Bopyridae | *Gyge* | *ovalis* | OP857530 | | | | | | NC037467 | KY038053 | | NC037467 | NC037467 |
| Cymothoida | Epicaridea: Bopyroidea | Bopyridae | *Argeia* | *pugettensis* | KF765770 | | | | | | MG753775 | | | MG753775 | MG753775 |
| Cymothoida | Epicaridea: Bopyroidea | Bopyridae | *Pleurocryptella* | *sp/fimbriata** | OL829930 | | | | | | MG729628 | MG729628 | | MG729628 | MG729628 |
| Cymothoida | Epicaridea: Bopyroidea | Ionidae | *Ione* | *cornuta* | KF765761 | | | | | | | | | | |
| Cymothoida | Epicaridea: Bopyroidea | Entoniscidae | *Portunion* | *conformis/sp/ sinensis** | KF765764 | | Table S1 | Table S1 | Table S1 | Table S1 | OL677861 | OL677861 | | OL677861 | MG936161 |
| Cymothoida | Epicaridea: Bopyroidea | Entoniscidae | *Cancrion* | *carolinus* | OQ171438 | | | | | | | | | | |

| | | | | | | | | | | | | |
|---|---|---|---|---|---|---|---|---|---|---|---|---|
| Cymothoida | Epicaridea: Cryptoniscoidea | Dajidae | *Zonophryxus* | *quinquedens* | DQ008451 | | | | | | | |
| Cymothoida | Epicaridea: Cryptoniscoidea | Dajidae | *Holophryxus* | *sp* | OL323109 | | | | | | | OK489803 |
| Cymothoida | Epicaridea: Cryptoniscoidea | Entophilidae | *Entophilus* | *mirabiledictu* | KF765763 | | | | | | | |
| Cymothoida | Epicaridea: Cryptoniscoidea | Cyproniscidae? | *Cryptoniscoidea* | *sp* | KF765772 | | | | | | | |
| Cymothoida | Epicaridea: Cryptoniscoidea | Cabiropidae?? | *Cymodoce* | *tattersalli* | AF255695 | | | | | | | |
| Phreatoicidea | | Amphisopodidae | *Eophreatoicus* | *sp* | KF260958 | | | | EU263213 | NC013976 | NC013976 | FJ790313 |
| Phreatoicidea | | Amphisopodidae | *Paramphisopus* | *palustris* | AY781425 | | | | AF259533 | AF259523 | | GQ926939 |
| Phreatoicidea | | Hypsimetopidae | *Pilbarophreatoicus /Andhracoides** | *shabuddin/sp** | KC771215 | | | | | DQ352854 | | OR653474 |
| Phreatoicidea | | Phreatoicidae | *Colubotelson* | *thomsoni/sp** | AF255703 | AF169711 | | | AF259531 | AF259524 | | AF255775 |
| Phreatoicidea | | Phreatoicopsidae | *Phreatoicopsis* | *raffae* | GQ302714 | | | | GQ302688 | | | GQ302698 |
| Microcerberidea | | Microcerberidae | *Coxicerberus* | *fukudai* | MG253041 | | | | | | | MF346640 |
| Asellota | Janiroidea | Acanthaspidiidae | *Ianthopsis* | *multispinosa* | EU414419 | | | | AY691342 | | | |
| Asellota | Janiroidea | Acanthaspidiidae | *Acanthaspidia* | *drygalskii* | EU414416 | | | | AY691369 | | | |
| Asellota | Janiroidea | Dendrotionidae | *Dendrotion* | *sp* | EU414423 | | | | CCZ1695-17 | | | CCZ1695-17 |
| Asellota | Janiroidea | Dendrotionidae | *Acanthomunna* | *spinipes* | EU414421 | | | | | | | |
| Asellota | Janiroidea | Dendrotionidae | *Dendromunna* | *sp* | EU414348 | | | | | | | |
| Asellota | Janiroidea | Desmosomatidae | *Mirabilicoxa* | *sp* | AY461461 | | | | MN015788 | | | KJ736035 |
| Asellota | Janiroidea | Desmosomatidae | *Chelator* | *insignis/sp** | AY461460 | | | | KJ630815 | | | KJ736036 |
| Asellota | Janiroidea | Desmosomatidae | *Eugerda* | *sp* | AY461463 | | | | MF325640 | | | MF325481 |
| Asellota | Janiroidea | Desmosomatidae | *Eugerdella* | *natator/huberti** | AY461462 | | | | HQ214679 | | | HQ214677 |
| Asellota | Janiroidea | Haploniscidae | *Antennuloniscus* | *sp* | EU414426 | | | | AY693397 | | | |
| Asellota | Janiroidea | Haploniscidae | *Chauliodoniscus* | *sp* | EU414427 | | | | MN550021 | | | MW066922 |
| Asellota | Janiroidea | Haploniscidae | *Haploniscus* | *rostratus/sp** | EU414429 | | | | AY693420 | | | JF283461 |
| Asellota | Janiroidea | Haploniscidae | *Mastigoniscus* | *polygomphios/sp** | EU414434 | | | | AY693399 | | | KJ736147 |
| Asellota | Janiroidea | Haplomunnidae | *Thylakogaster* | *sp* | EU414424 | | | | CCZ1667-17 | | | KJ736121 |
| Asellota | Janiroidea | Ischnomesidae | *Haplomesus* | *sp* | AY461474 | | | | | | | KJ736063 |
| Asellota | Janiroidea | Ischnomesidae | *Ischnomesus* | *sp/bispinosus** | EU414435 | OQ071323 | | | DISCO091 | | | KJ736062 |
| Asellota | Janiroidea | Ischnomesidae | *Stylomesus* | *sp* | EU414436 | | | | | | | MN311579 |
| Asellota | Janiroidea | Janirellidae | *Janirella* | *sp* | AY461475 | | | | | | | LC535322 |
| Asellota | Janiroidea | Janiridae | *Ianiropsis* | *epilittoralis* | EF682260 | EF682305 | | | AF260859 | | | EF682303 |
| Asellota | Janiroidea | Janiridae | *Janira* | *maculosa/sp** | AF255700 | | | | AJ388079 | GU130255 | GU130255 | GU130255 |
| Asellota | Janiroidea | Janiridae | *Jaera* | *albifrons* | AF279609§ | ON598986 | MH880902 | | AJ388078 | | | ON601009 |
| Asellota | Janiroidea | Janiridae | *Neojaera* | *sp* | EU414439 | | | | | | | |
| Asellota | Janiroidea | Janiridae | *Iathrippa* | *trilobatus* | AF279606 | | | | | | | |
| Asellota | Janiroidea | Janiridae | *Iais* | *pubescens* | EU414437 | | | | | | | |
| Asellota | Janiroidea | Joeropsidae | *Joeropsis* | *coralicola/dubia** | AF279608 | | | | AF260860 | | | KX608735 |
| Asellota | Janiroidea | Macrostylidae | *Macrostylis* | *sp* | AY461477 | | | | LT909395 | JX260275 | | CCZ1404-17 |
| Asellota | Janiroidea | Mesosignidae | *Mesosignum* | *weddellensis/sp** | EU414443 | | | | CCZ1083-17 | | | CCZ1079-17 |
| Asellota | Janiroidea | Munnidae | *Uromunna* | *nana* | KX467630 | | | | | | | KJ736157 |
| Asellota | Janiroidea | Munnopsidae; Acanthocopinae | *Acanthocope* | *galathea/sp** | EF682241 | EF682336 | | | MG722005 | | | MG721975 |
| Asellota | Janiroidea | Munnopsidae; Betamorphinae | *Betamorpha* | *africana/fusiformis** | EF682248 | EF682331 | | | EF116541 | | | KJ736131 |
| Asellota | Janiroidea | Munnopsidae; Eurycopinae | *Eurycope* | *glabra/hanseni** | EF682255 | EF682329 | | MH056537 | MH056219 | | | MH056598 |
| Asellota | Janiroidea | Munnopsidae; Eurycopinae | *Dubinectes* | *acutitelson* | EF682251 | EF682330 | | | | | | EF682294 |
| Asellota | Janiroidea | Munnopsidae; Ilyarachninae | *Ilyarachna* | *triangulata/sp** | EF682244 | EF682333 | | | MN550437 | | | EF682299 |

| | | | | | | | | | | | | | | | |
|---|---|---|---|---|---|---|---|---|---|---|---|---|---|---|---|
| Asellota | Janiroidea | Munnopsidae; Ilyarachninae | *Notopais* | *magnifica/sp** | EF682249 | | Table S1 | | | Table S1 | OL661186 | | | | OL661186 |
| Asellota | Janiroidea | Munnopsidae; Lipomerinae | *Mimocopelates* | *sp* | EU414460 | EF682328 | | | | | | | | | EF682297 |
| Asellota | Janiroidea | Munnopsidae; Munnopsinae | *Munnopsis* | *abyssalis/sp** | EF682225 | EF682313 | | | | | MN550102 | | | | HQ919177 |
| Asellota | Janiroidea | Munnopsidae; Bathyopsurinae | *Paropsurus* | *giganteus* | EF682253 | EF682339 | | | | | | | | | EF682287 |
| Asellota | Janiroidea | Munnopsidae; Storthyngurinae | *Storthyngurella* | *menziesi* | EU414464 | | | | | | | | | | MW072543 |
| Asellota | Janiroidea | Nannoniscidae | *Austroniscus* | *sp* | EU414469 | | | | | | MN015685 | | | | MZ151074 |
| Asellota | Janiroidea | Nannoniscidae | *Nannoniscus* | *sp* | EU414470 | | | | | | MF325664 | | | | KJ736099 |
| Asellota | Janiroidea | incertae | *Xostylus* | *sp* | EU414471 | | | | | | | | | | KJ736158 |
| Asellota | Stenetrioidea | Stenetriidae | *Tenupedunculus* | *sp* | EU414473 | | | | | | | | | | |
| Asellota | Stenetrioidea | Stenetriidae | *Stenetriid* | *sp* | AY461453 | | | | | | | | | | OGL455-11 |
| Asellota | Aselloidea | Stenasellidae | *Stenasellus* | *racovitzai/virei** | AF496663 | JQ922016 | | | | | JQ921837 | | | | JQ921624 |
| Asellota | Aselloidea | Asellidae | *Asellus* | *aquaticus* | AF255701 | MN174846 | MH880900 | | MN234323 | AJ238321 | MG205328 | MG205174 | | GU130252 | GU130252 |
| Asellota | Aselloidea | Asellidae | *Lirceus* | *fontinalis/culveri/sp** | AF255702 | | Table S1 | Table S1 | Table S1 | Table S1 | OX383335 | | | | OQ918606 |
| Asellota | Aselloidea | Asellidae | *Proasellus* | *slavus/racovitzai/ walteri** | AF496662 | JQ921975 | Table S1 | Table S1 | Table S1 | Table S1 | KC610209 | LR536620 | | LR536620 | JQ921389 |
| Asellota | Aselloidea | Asellidae | *Caecidotea* | *racovitzai/kenki/sp** | AY781426 | JQ921877 | | | | | KC610276 | AF259529 | | | JQ921575 |
| Amphipoda | | | *Monoculodes* | *packardi* | DQ378015 | | | | | | MN228726 | | | | |
| Amphipoda | | | *Paroediceros* | *propinquus* | AF419231 | | | | | | AMPIV086-17 | | | | AMPIV087-17 |
| Amphipoda | | | *Arrhis* | *phyllonyx* | AF419235 | | | | | | AMPIV088-17 | | | | AMPIV088-17 |
| Amphipoda | | | *Gammarus* | *duebeni/electrus** | AF356545 | | Table S1 | | | Table S1 | JN704067 | | NC017760 | | JN704067 |
| Amphipoda | | | *Astyra* | *antarctica* | DQ377999 | | | | | KF484717 | KF430288 | | | | MG264764 |
| Amphipoda | | | *Talorchestia* | *spinipalma* | MG655746§ | | | | | MG655998 | MG655686 | | | | MG655889 |
| Stygiomysida | | | *Stygiomysis* | *cokei* | AM422478 | | | | | | | | | | |
| Spelaeogriphacea | | | *Spelaeogriphus* | *lepidops* | AY781414§ | | | | | | | | | | |
| Tanaidacea | | | *Apseudes* | *bermudeus/latreillei** | GQ175865 | | Table S1 | | | Table S1 | AJ388110 | | | | MW284787 |
| Tanaidacea | | | *Tanais* | *dulongii* | AY781428 | | | | | KF928332 | | | | | HM016204 |
| Cumacea | | Nannastacidae | *Cumella* | *sp* | MK635505 | | Table S1 | | | Table S1 | MK613880 | | OL841458 | | MK757519 |
| Cumacea | | Diastylidae | *Diastylis* | *sculpta/cornuta* | AY781431 | | Table S1 | | | Table S1 | U81512 | | | | AF137510 |

**Supplementary Table S4.** Marker-gene dataset parameters, including base composition analysis in *PAUP** (a significant *p*-value indicates that the partition failed the base composition test, requiring RY-recoding. RY-recoded partitions which failed a base composition test were excluded)*,* as well as best model selected for phylogenetic analysis in *IQ-TREE* and molecular dating in *BEAST*.

| Genome | Gene | Partition | Length (base pairs) | 4-nt coding *(p-value)* | RY-recoded *(p-value)* | ML model selected | BEAST model |
|---|---|---|---|---|---|---|---|
| **Nuclear** | **18S** | | 1990 | 1.000 | | SYM+R6 | SYM+G |
| | **28S** | | 320 | 1.000 | | SYM+I+G4 | SYM+I+G |
| | **NAK** | 1 | 209 | 1.000 | | SYM+R3 | SYM+G |
| | | 2 | 208 | 1.000 | | TNe+R2 | TNe+G |
| | | 3 | 208 | **0.000** | 1.000 | TVM+F+I | HKY+I |
| | **PEPCK** | 1 | 169 | 1.000 | | TPM2+I+G4 | TPM2+I+G |
| | | 2 | 168 | 1.000 | | SYM+I+I+R2 | SYM+I+G |
| | | 3 | 168 | **0.000** | 1.000 | TPM2+I+G4 | HKY+I+G |
| | **IGFPB** | 1 | 265 | 1.000 | | GTR+F+I+G4 | GTR+F+I+G |
| | | 2 | 265 | 1.000 | | TPM2+G4 | TPM2+G |
| | | 3 | 265 | **0.000** | 1.000 | TVMe+I | HKY+G |
| | **H3** | 1 | 124 | 1.000 | | SYM+I+G4 | SYM+I+G |
| | | 2 | 124 | 1.000 | | JC | JC |
| | | 3 | 123 | **0.000** | 1.000 | TIM2e+I+G4 | HKY+I+G |
| **Mitochondrial** | **12S** | | 595 | 0.384 | | GTR+F+G4 | GTR+F+G |
| | **16S** | | 438 | 0.248 | | GTR+F+R6 | GTR+F+G |
| | **COI** | 1 | 219 | 0.979 | | GTR+F+R5 | GTR+F+G |
| | | 2 | 219 | 1.000 | | GTR+F+R4 | GTR+F+G |
| | | 3 | 219 | **0.000** | **0.000** | | |
| | **ND4** | 1 | 312 | **0.000** | 1.000 | SYM+G4 | HKY+G |
| | | 2 | 312 | **0.000** | 1.000 | GTR+F+G4 | HKY+F+G |
| | | 3 | 312 | **0.000** | **0.000** | | |
| | **CYTB** | 1 | 308 | **0.000** | 0.927 | TVMe+G4 | HKY+G |
| | | 2 | 308 | 0.582 | | TVM+F+G4 | TVM+F |
| | | 3 | 308 | **0.000** | **0.000** | | |

**Supplementary Table S5.** Fossil calibrations for the *BEAST* analysis. A hard upper bound (maximum constraint) of 521 mya was set for the root, which marks the appearance of crown group arthropods in the fossil record (1). A soft maximum constraint (upper 97.5% of log normal distribution) of 521 mya was applied to internal fossil constraints 2-6, and of 359.3 mya to constraints 7-16, which marks the appearance of the first peracarid Crustacea in the fossil record (2,3).

| | **Taxa** | **BEAST parameters** | **Divergence** | **Lower boundary** | **Fossil taxa** | **Date and Locality Information** | **Refs** |
|---|---|---|---|---|---|---|---|
| 1 | All taxa | uniformPrior lower="359.3" upper="521.0" | Root of tree | 359.3 mya | *Tealliocaris walloniensis* | Late Devonian (Famennian); Bois des Mouches Formation of Belgium | 2,3 |
| 2 | Tanaidacea | logNormalPrior mean="158.3" stdev="79.2" offset="163.5" | Base of Tanaidacea | 163.5 mya | *Palaeotanais quenstedti* | Middle Jurassic of southern Germany | 4 |
| 3 | Cumacea | logNormalPrior mean="191.0" stdev="95.45" offset="90.0" | Base of Cumacea | 90 mya | *Eobodotria muisca* | Mid-Cretaceous Lagerstätte of Colombia | 5 |
| 4 | Isopoda | logNormalPrior mean="95.1" stdev="47.3" offset="307.0" | Split between Phreatoicidea & Asellota | 307 mya | *Hesslerella shermani* | Carboniferous (Pennsylvanian); Mazon Creek, Francis Creek Shale, Braidwood, Illinois, USA | 6 |
| 5 | Phreatoicidea: Amphisopodidae | logNormalPrior mean="89.4" stdev="44.65" offset="242.0" | Split between Amphisopodidae & other Phreatoicids | 242 mya | *Protamphisopus wianamattensis* | Middle Triassic (Anisian) Ashfield Shale in the Sydney Basin, Australia | 7 |
| 6 | Asellota: Janiroidea | logNormalPrior mean="104.2" stdev="52.1" offset="208.5" | Base of Janiroidea | 208.5 mya | *Fornicaris calligarisi* | Triassic, (Norian); Dolomia di Forni Formation, Forni di Sotto, NE Italy, | 8 |
| 7 | Asellota: Stenetrioidea | logNormalPrior mean="155" stdev="77.45" offset="93.9" | Split within Stenetriidae | 93.9 mya | *Eostenetrium guerangeri* | Upper Cretaceous (Cenomanian stratotype); Le Mans, Western Paris Basin, France | 9 |
| 8 | Epicaridea: Bopyroidea | logNormalPrior mean="95.0" stdev="47.45" offset="145.0" | Split between Bopyroidea & Cryptoniscoidea | 145 mya | *Mesogalathea striata* | Carapace swelling indicating bopyroid infection; Jurassic (Kimmeridgian); Czechia & Austria | 10 |
| 9 | Cymothooidea | logNormalPrior mean="92.4" stdev="46.15" offset="150.8" | Base of Cymothooidea (Cirolanidae) | 150.8 mya | *Brunnaega roeperi* | Late Jurassic, Plattenkalk of Brunn, southern Germany | 11 |
| 10 | Sphaeromatidea + Valvifera | logNormalPrior mean="66.8" stdev="33.4" offset="208.5" | Split between Valvifera & Sphaeromatidea | 208.5 mya | *Triassphaeroma magnificum* | Triassic (Norian); Southern Calcareous Alps, Calcare di Zorzino, Northern Italy, | 12 |
| 11 | Sphaeromatidea | logNormalPrior mean="95.0" stdev="47.45" offset="145.0" | Split between Seroloidea & Sphaeromatoidea | 145 mya | *Schweglerella strobli* | Late Jurassic (Early Tithonian); Plattenkalk of Solnhofen, southern Germany | 13 |
| 12 | Valvifera: Chaetiliidae | logNormalPrior mean="94.7" stdev="47.1" offset="93.9" | Split between Chaetiliidae & Antarcturidae | 93.9 mya | *Protochaetilia delaunayi* | Upper Cretaceous (Cenomanian stratotype); Le Mans, Western Paris Basin, France | 9 |
| 13 | Valvifera: Idoteidae | logNormalPrior mean="94.7" stdev="47.1" offset="93.9" | Split between Idoteidae & Holognathidae | 93.9 mya | *Mesozoidotea gazonfierensis* | Upper Cretaceous (Cenomanian stratotype); Le Mans, Western Paris Basin, France | 9 |
| 14 | Oniscidea: Diplocheta | logNormalPrior mean="89.5" stdev="44.73" offset="105.0" | Base of Diplocheta | 105.0 mya | *Eoligiiscus tarraconensis* | Cretaceous, (Albian) Amber from Peñacerrada I outcrop, Burgos Provinces, Northern Spain | 14 |
| 15 | Oniscidea: Synocheta | logNormalPrior mean="89.5" stdev="44.73" offset="105.0" | Base of Synocheta | 105.0 mya | *Autrigoniscus resinicola* | Cretaceous, (Albian) Amber from Peñacerrada I outcrop, Burgos Provinces, Northern Spain | 14 |
| 16 | Oniscidea: Crinocheta | logNormalPrior mean="89.5" stdev="44.73" offset="105.0" | Base of Crinocheta | 105.0 mya | *Heraclitus helenae* | Cretaceous, (Albian) Amber from Peñacerrada I outcrop, Burgos Provinces, Northern Spain | 14 |

**Table S6.** Results for tests of topology (see table footnote), performed in *IQ-TREE*. There were 11 alternative hypotheses tested against the ML tree, illustrated in Supplementary Figure S3. The only topology which could not be rejected was the topology recovered by the *ASTRAL* summary tree, given in Supplementary Figure S5, where Limoriidea was recovered as sister to Valvifera/Sphaermatoidea.

| Hypothesis | Tree | logL | deltaL | bp-RELL | Sig? | p-KH | Sig? | p-SH | Sig? | p-WKH | Sig? | p-WSH | Sig? | c-ELW | Sig? | p-AU | Sig? |
|---|---|---|---|---|---|---|---|---|---|---|---|---|---|---|---|---|---|
| (a) Erhard | 1 | -5841104.345 | 169.8 | 0.000 | - | 0.000 | - | 0.159 | - | 0.000 | - | 0.000 | - | 1.33e-17 | - | 1e-06 | - |
| (b) MarineD | 2 | -5845214.913 | 4280.4 | 0.000 | - | 0.000 | - | 0.000 | - | 0.000 | - | 0.000 | - | 0.000 | - | 4.82e-44 | - |
| (c) MarineT | 3 | -5843364.902 | 2430.4 | 0.000 | - | 0.000 | - | 0.000 | - | 0.000 | - | 0.000 | - | 0.000 | - | 7.57e-79 | - |
| (d) MarineDT | 4 | -5845212.724 | 4278.2 | 0.000 | - | 0.000 | - | 0.000 | - | 0.000 | - | 0.000 | - | 0.000 | - | 7.96e-40 | - |
| (e) CLimn | 5 | -5840957.389 | 22.8 | 0.0068 | - | 0.0073 | - | 0.797 | - | 0.0073 | - | 0.0449 | - | 0.0079 | - | 0.00519 | - |
| (f) ELimn | 6 | -5841336.442 | 401.9 | 0.000 | - | 0.000 | - | 0.0041 | - | 0.000 | - | 0.000 | - | 8.45e-105 | - | 5.19e-67 | - |
| **(g) VSLimn** | **7** | **-5840934.549** | **0** | **0.993** | + | **0.993** | + | **1.000** | + | **0.993** | + | **1.000** | + | **0.992** | + | **0.995** | + |
| (h) VSLimn_Erhard | 8 | -5841142.287 | 207.7 | 0.000 | - | 0.000 | - | 0.0944 | - | 0.000 | - | 0.000 | - | 8.14e-44 | - | 1.35e-31 | - |
| (i) VSLimn_MarineD | 9 | -5845247.321 | 4312.8 | 0.000 | - | 0.000 | - | 0.000 | - | 0.000 | - | 0.000 | - | 0.000 | - | 4.94e-45 | - |
| (j) VSLimn_MarineT | 10 | -5843405.107 | 2470.6 | 0.000 | - | 0.000 | - | 0.000 | - | 0.000 | - | 0.000 | - | 0.000 | - | 4.99e-52 | - |
| (k) VSLimn_MarineDT | 11 | -5845247.549 | 4313.0 | 0.000 | - | 0.000 | - | 0.000 | - | 0.000 | - | 0.000 | - | 0.000 | - | 3.3e-48 | - |

**deltaL**: logL difference from the maximal logl in the set.
**bp-RELL**: bootstrap proportion using RELL method (Kishino et al. 1990).
**p-KH**: p-value of one sided Kishino-Hasegawa test (1989).
**p-SH**: p-value of Shimodaira-Hasegawa test (2000).
**p-WKH**: p-value of weighted KH test.
**p-WSH**: p-value of weighted SH test.
**c-ELW**: Expected Likelihood Weight (Strimmer & Rambaut 2002).
**p-AU**: p-value of approximately unbiased (AU) test (Shimodaira, 2002).

Plus-signs denote the 95% confidence sets.
Minus signs denote significant exclusion.
All tests performed 10000 resamplings using the RELL method.
