## Supplementary Figures for "Phylogenomics supports a single origin of terrestriality in Isopods"

**Supplementary Figure S1.** Pictures of representative species in each major group of Isopoda

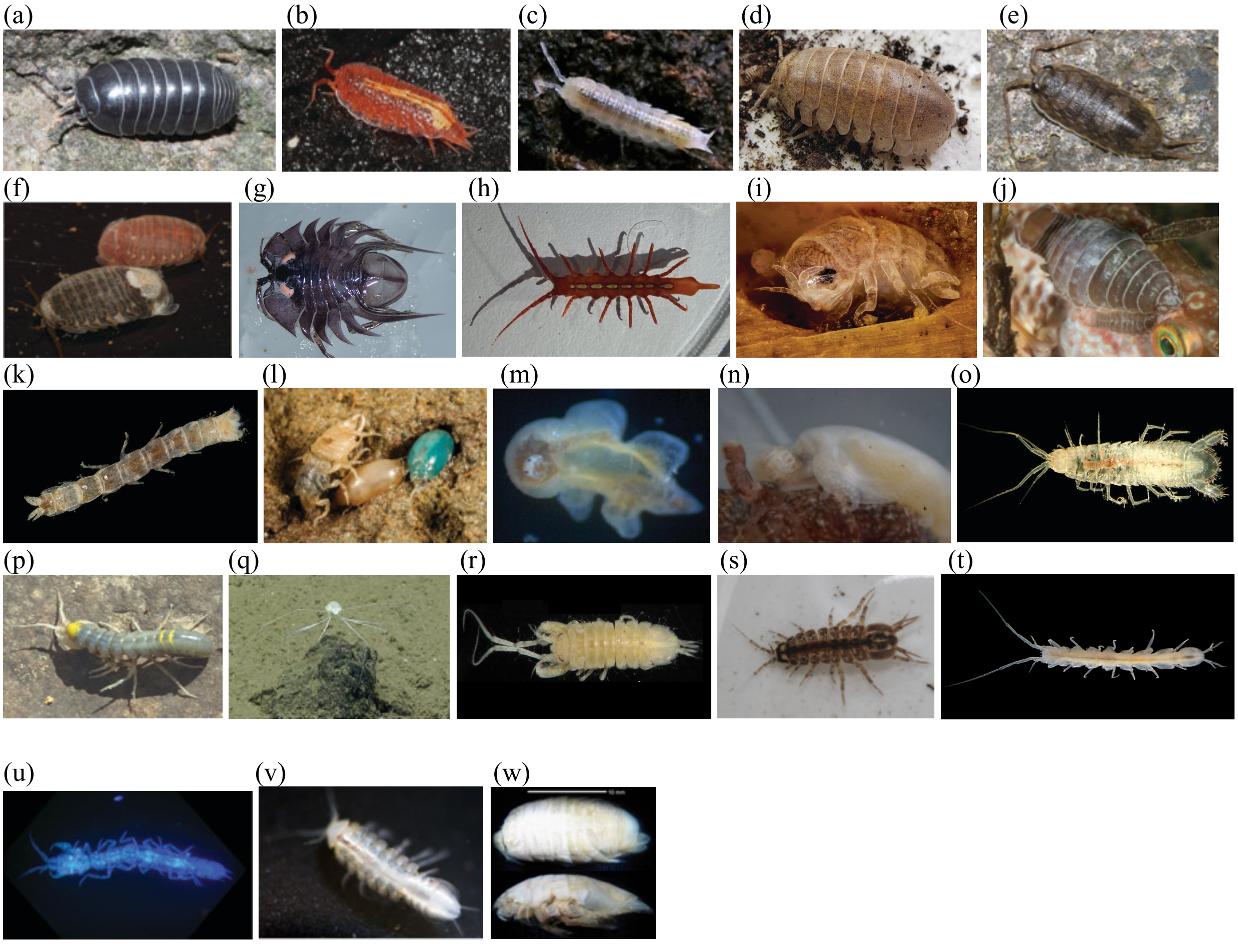


**Isopod Suborders:**

**Oniscidea**: (a) **Crinocheta**, *Armadillidium vulgare* © W.Maguire, British Myriapod & Isopod Group (BMIG)
(b) **Synocheta**, *Androniscus dentiger* © W.Maguire, BMIG, https://bmig.org.uk/
(c) **Microcheta**, *Mesoniscus graniger* © L.Kováč & P.Luptacik. Reprint from Smrz J, Kovac L, Mikes J, Sustr V, Lukesova A, Tajovsky K, Novakova A, Reznakova P (2015) Food sources of selected terrestrial cave arthropods. *Subterranean Biology* 16: 37-46. doi: 10.3897/subtbiol.16.8609
(d) **Tylida**, *Helleria brevicornis* © T.Hughes, BMIG.
(e) **Diplocheta**, *Ligia oceanica* © W.Maguire, BMIG.

**Sphaeromatidea**: (f) **Sphaeromatoidea**, *Sphaeroma serratum* © W.Maguire, BMIG.
(g) **Seroloidea**, *Brucerolis sp* © A.Hosie, Western Australian Museum, Perth.

**Valvifera**: (h) *Stenosoma lancifer* © J.Thomas Thorpe, Darwin Tree of Life Project (DToL), Wellcome Sanger Institute.

**Limnoriidea:** (i) *Limnoria quadripuncata* © S.Trewhella.

**Cymothoida:** (j) **Cymothooidea**, *Anilocra sp* © S.Trewhella.
(k) **Anthuroidea**, *Cyathura carinata* © S.Trewhella.
(l) **Gnathiidea**, *Gnathia maxillaris* (male and two females) © S.Trewhella
(m) **Cryptoniscoidea**, *Hemioniscus balani* © P.Adkins, DToL, Marine Biological Association, Plymouth.
(n) **Bopyroidea,** *Athelges paguri* © J.Thomas Thorpe, DToL, Wellcome Sanger Institute, Cambridge.

**Tainisopidea**: (o) *Tainisopus sp* © G.D.F.Wilson.

**Phreatoicidea**: (p) *Phreatoicus sp* © G.D.F.Wilson.

**Asellota**: (q) **Janiroidea**, *Munnopsidae sp.* © NOAA.
(r) **Stenetrioidea**, *Stenetrium sp.* © G.D.F.Wilson.
(s) **Aselloidea**, *Asellus aquaticus* © W.Maguire, BMIG.
(t) **Gnathostenetroidoidea**, *Wiyufiloides osornoensis* © J.Perez-Schultheiss, G.D.F.Wilson, reprint from Perez-Schultheiss J, & Wilson GDF. (2021) A new genus and species of groundwater isopod of the family Protojaniridae (Isopoda: Asellota: Gnathostenetroidoidea) from southern Chile. Zootaxa, 4966(5):550–562, doi:10.11646/zootaxa.4966.5.4

**Microcerberidea**: (u) *Texicerberus sp.* © Benjamin Schwartz, Texas State University and Edwards Aquifer Research & Data Centre, https://www.eardc.txst.edu

**Calabozoidea**: (v) *Pongycarcinia xiphidiourus.* Reprint from: Prevorčnik, Simona, Ferreira, Rodrigo Lopes, Sket, Boris (2012): Brasileirinidae, a new isopod family (Crustacea: Isopoda) from the cave in Bahia (Brazil) with a discussion on its taxonomic position. Zootaxa 3452: 47-65, doi: 10.5281/zenodo.211428

**Phoratopidea**: (w) *Phoratopus remex,* © G.D.F.Wilson.

**Supplementary Figure S2.** Previous hypotheses of Isopod relationships

Relationships based on morphology:

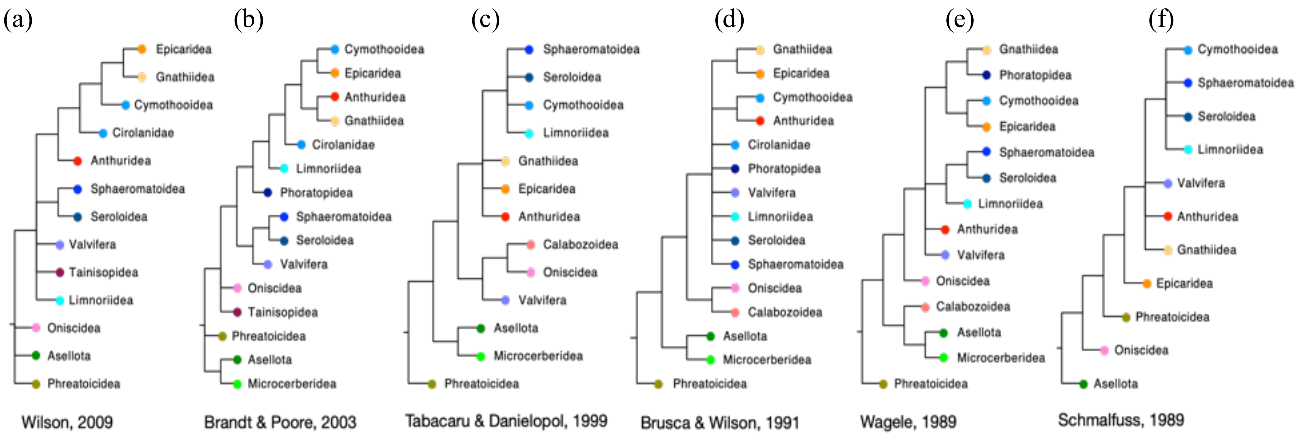


Relationships based on molecular data:

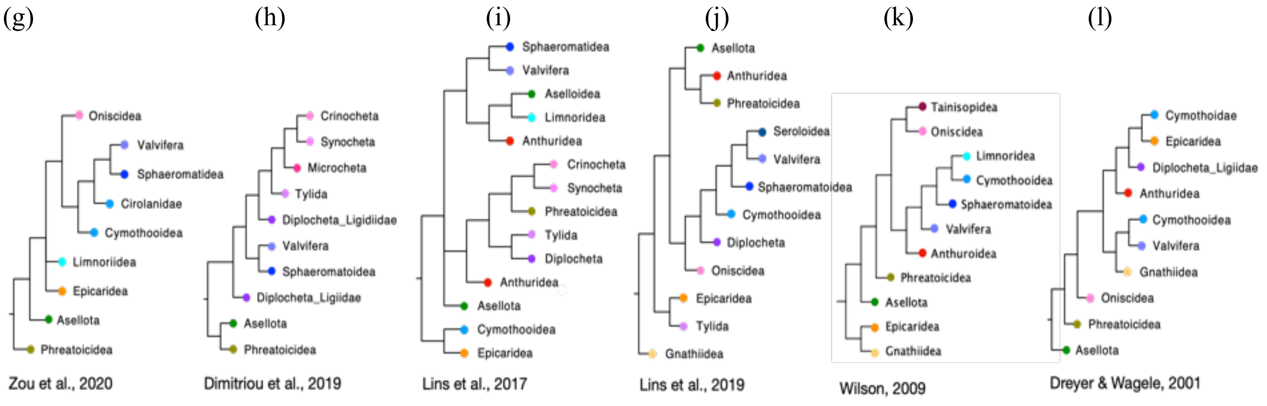


Relationships within Oniscidea based on morphology:

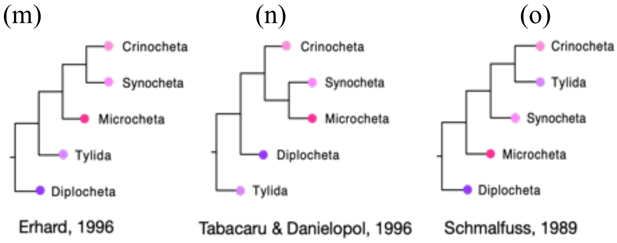

**Supplementary Figure S3.** Alternative hypotheses for tests of topology, performed in *IQ-TREE*. Corresponding results for this analysis are in Supplementary Table S6.

(a) Erhard

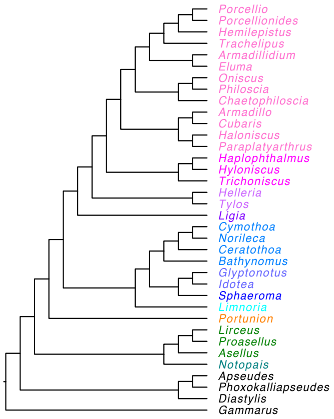


(b) Marine Diplocheta (MarineD)

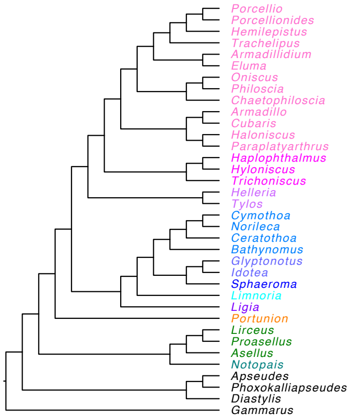


(c) Marine Tylida (MarineT)

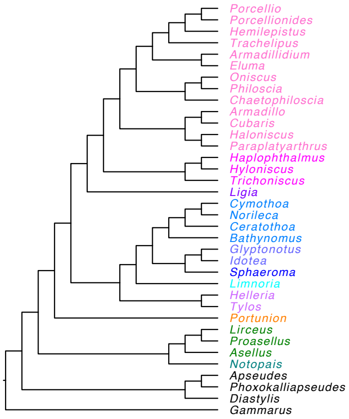


(d) Marine Tylida and Diplocheta (MarineDT)

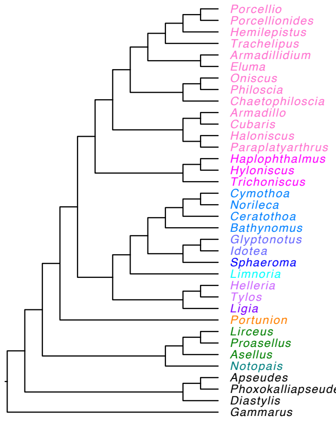


(e) Cymothoida-Limnoriidea (CLimn)

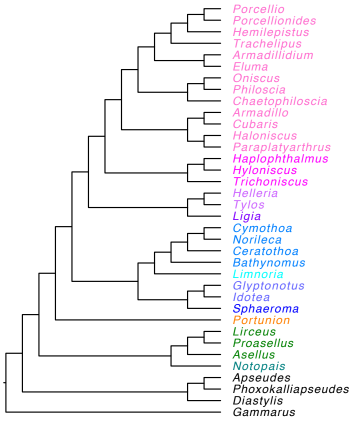


(f) Epicaridea-Limnoriidea (ELimn)

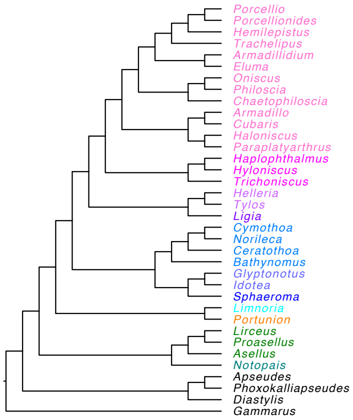


(g) Valvifera/Sphaeromatoidea-Limnoriidea (VSLimn)

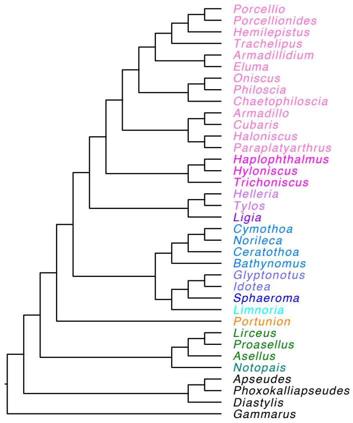


(h) VSLimn_Erhard

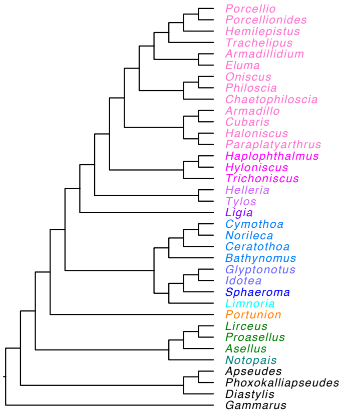


(i) VSLimn_MarineD

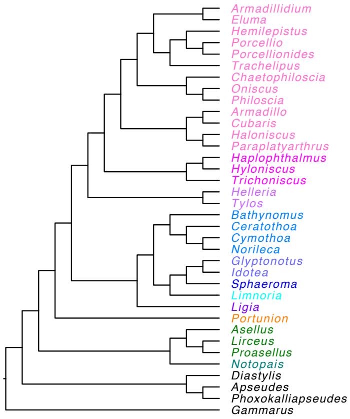


(j) VSLimn_MarineT

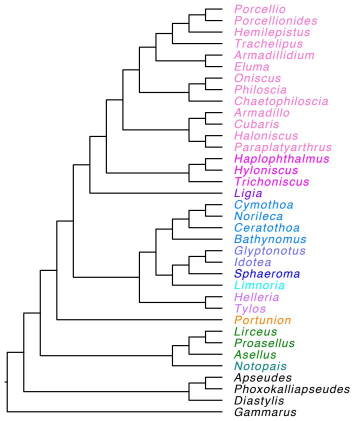


(k) VSLimn_MarineDT

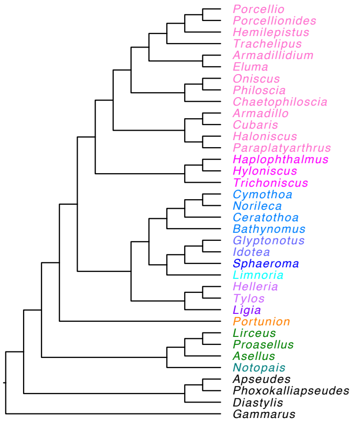

**Supplementary Figure S4 (a).** Dated phylogeny of *BEAST* analysis of marker-gene dataset, with Tainisopidea excluded, and nodes for clade 'CLVS' and 'CLVS'+Oniscidea unconstrained, to determine placement of Anthuroidea. Branch lengths in millions of years, values at nodes indicate posterior probability.

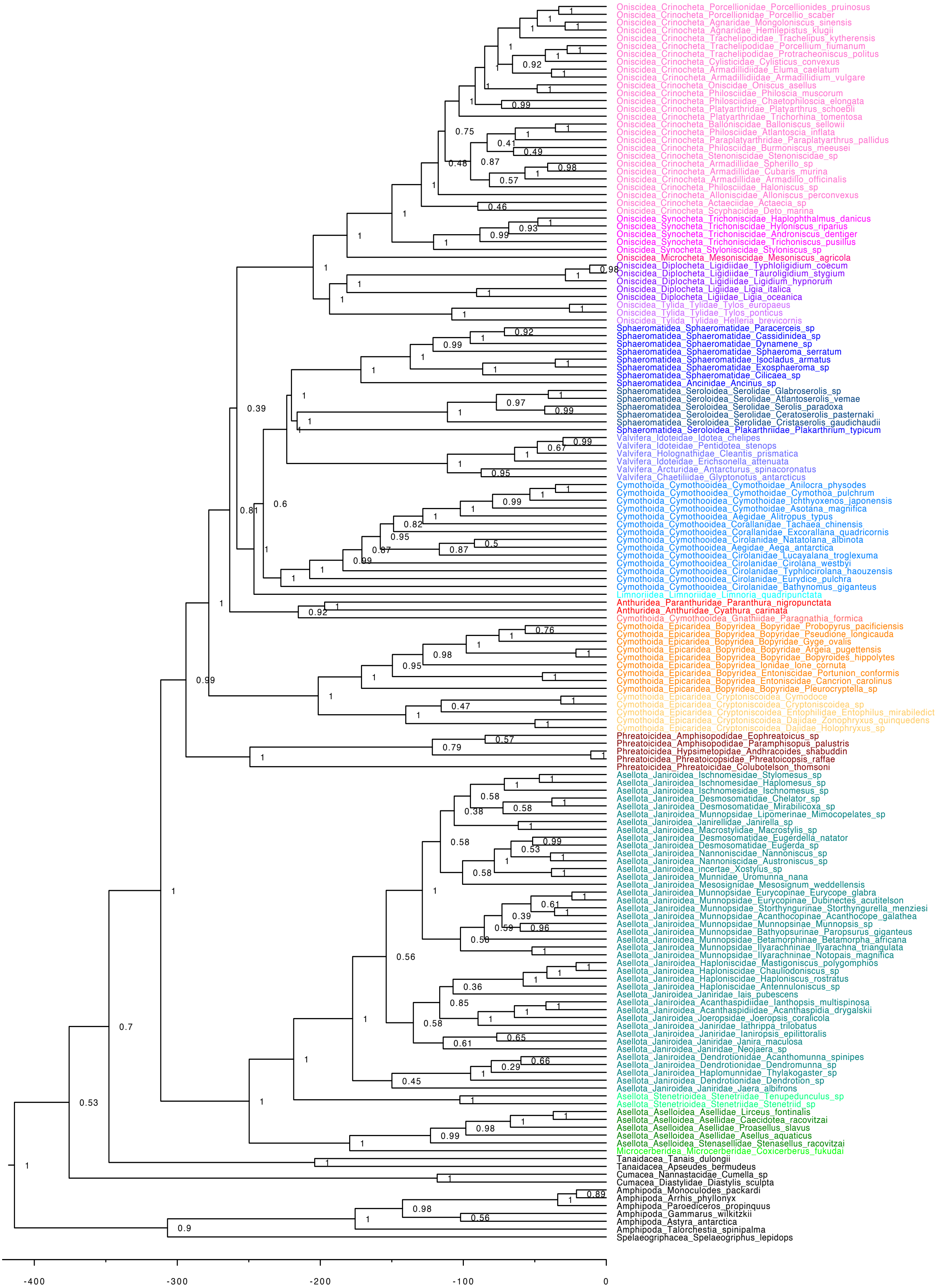

**Supplementary Figure S4 (b).** Dated phylogeny of *BEAST* analysis of marker-gene dataset, with Anthuroidea excluded, and nodes for clade 'CLVS' and 'CLVS'+Oniscidea unconstrained, to determine placement of Tainisopidea. Branch lengths in millions of years, values at nodes indicate posterior probability.

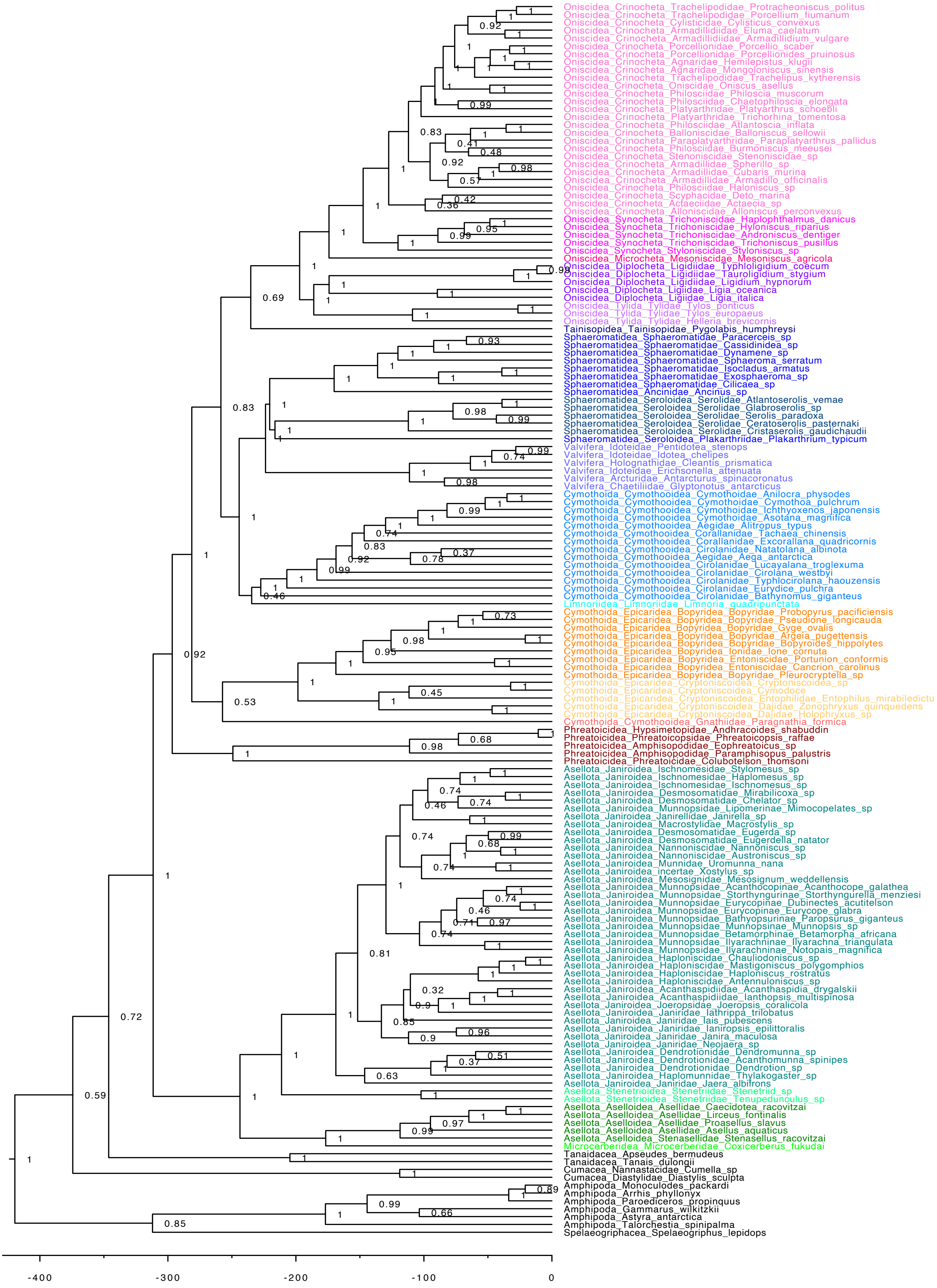

**Supplementary Figure S5.** ML phylogeny inferred from supermatrix of 960 single-copy orthologues, aligned with *FSA* and trimmed with *Trimal* (phylogenomic dataset) in *IQ-TREE*. All bootstrap values equal 100%. Identical topologies were produced by all four versions of the amino-acid datasets and the 1st and 2nd codon positions of the nucleotide dataset. Branch lengths are in substitution per site.

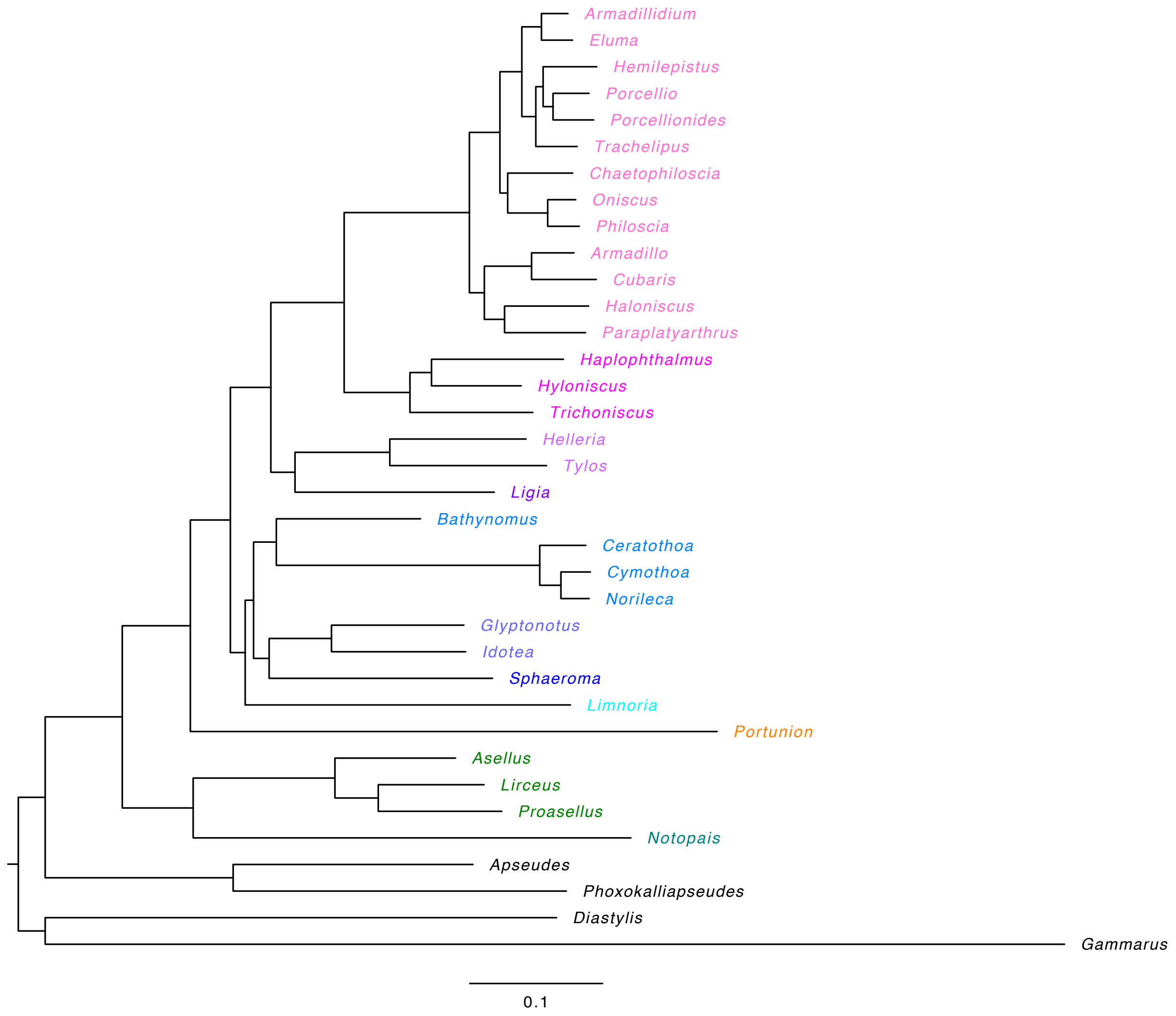

**Supplemental Figure S6.** Summary of individual gene trees, produced in *ASTRAL*. All versions of the dataset produced an identical topology. Nodal values indicate bootstrap support for node. Branch lengths are measured in coalescent units.

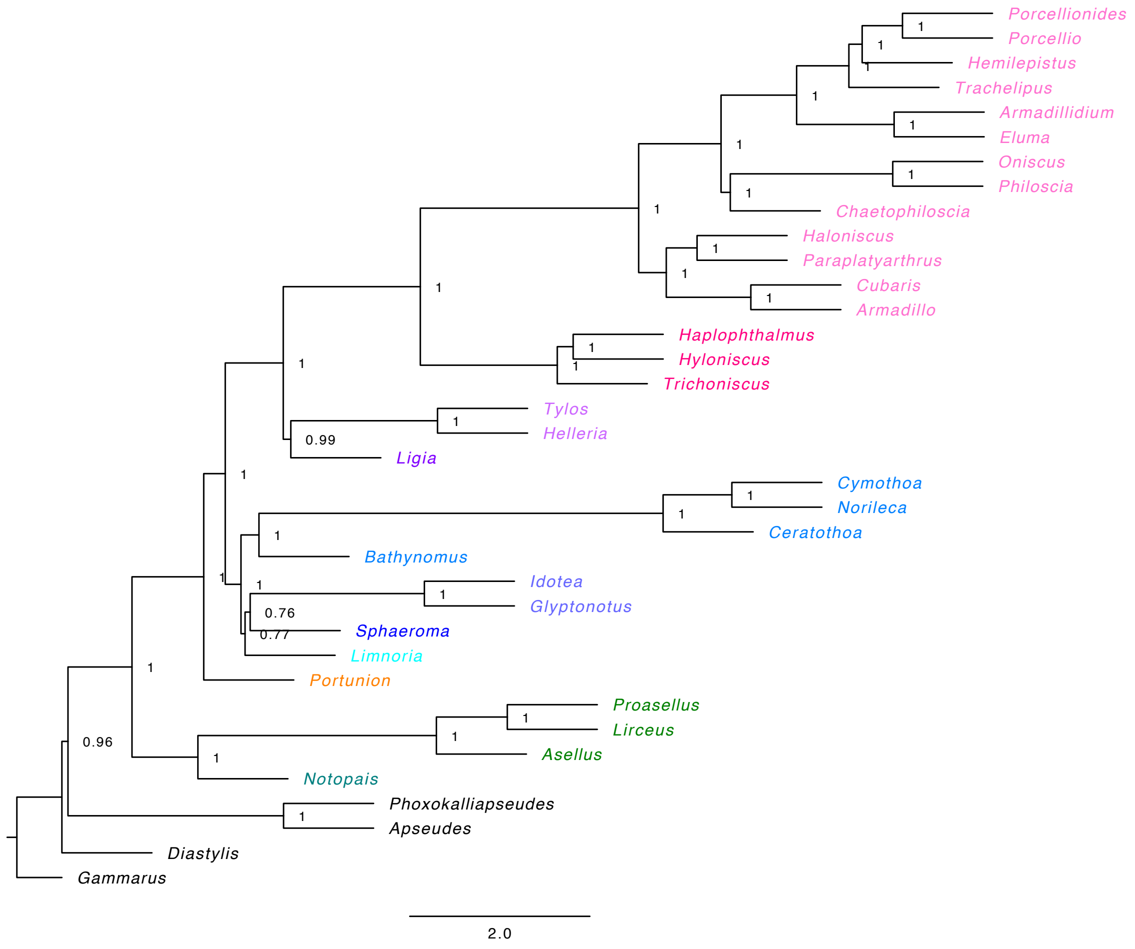

**Supplementary Figure S7:** Relationship between Gene Concordance Factors and gene length. Scatterplot of gene concordance factors for each node of interest, recalculated across ten different iterations of the analysis, removing the shortest 10% of genes each time. Nodes are (A) Asellota, (B) Oniscidea and (C) 'CLVS'. Plots indicate a reduction in the proportion of polyphyletic relationships at each node as the shortest genes are removed. Data and code to produce plots are available in the Github repository https://github.com/jessthomasthorpe/Isopod_Phylogenomics_MS.

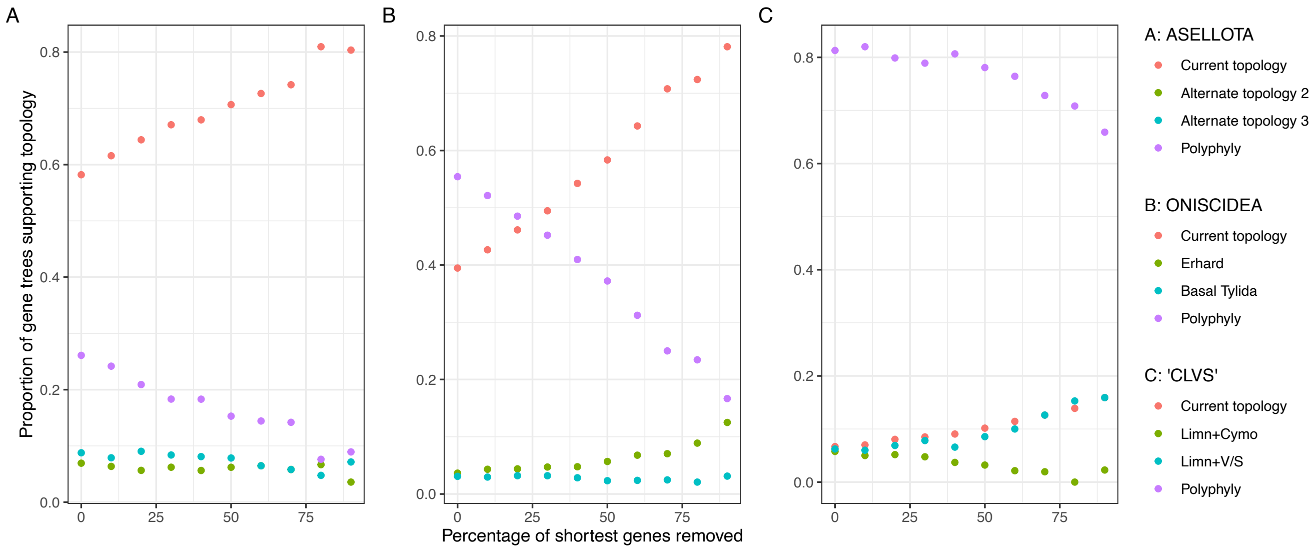


.

**Supplementary Figure S8:** Phylogeny from ML analysis in *IQ-TREE* of supermatrix of 11 marker-genes, values at nodes indicate bootstrap support. Branch lengths in substitutions per site.

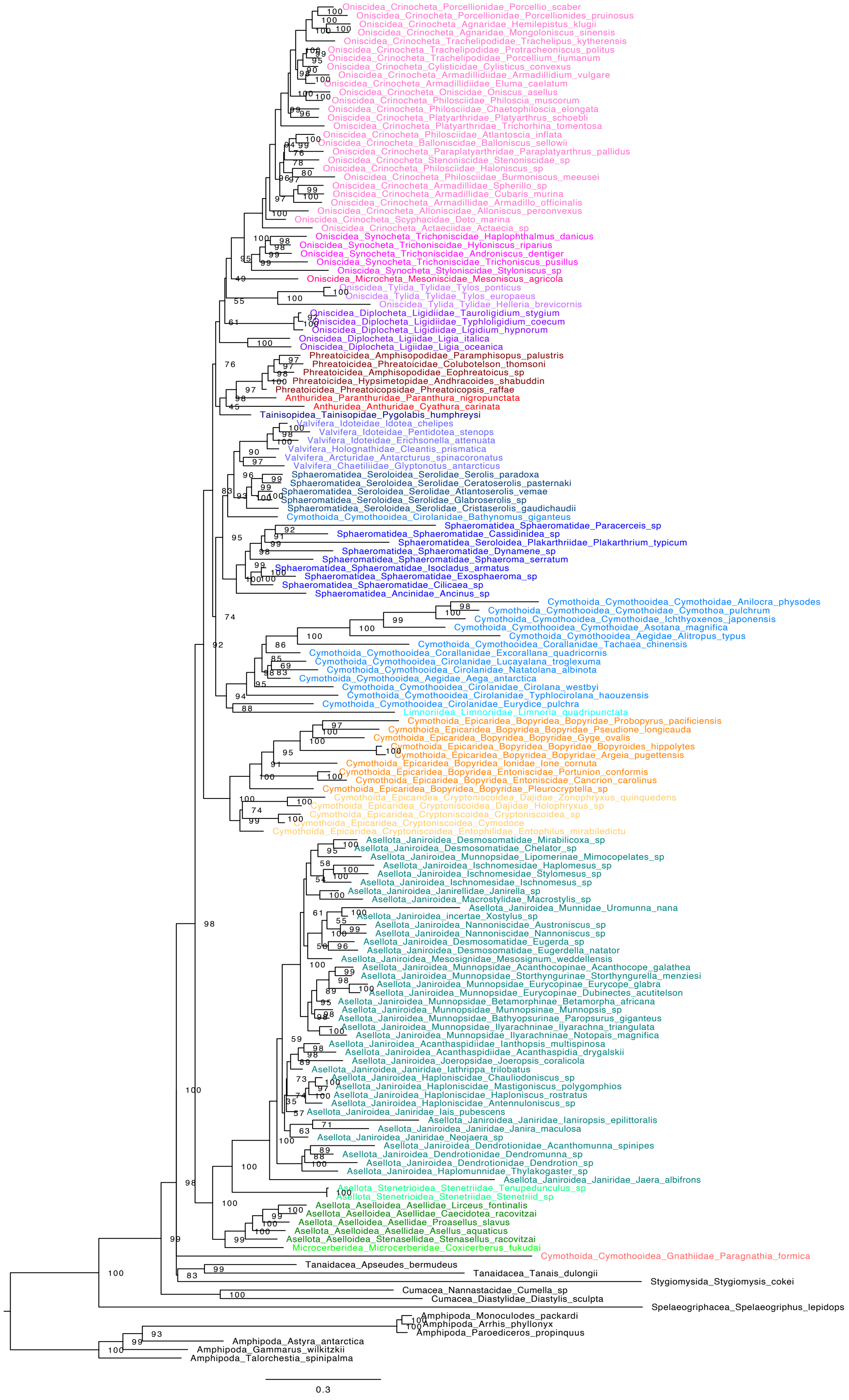

**Supplementary Figure S9:** Phylogeny from ML analysis in *IQ-TREE* of marker-gene dataset 18S only. Branch lengths in substitutions per site.

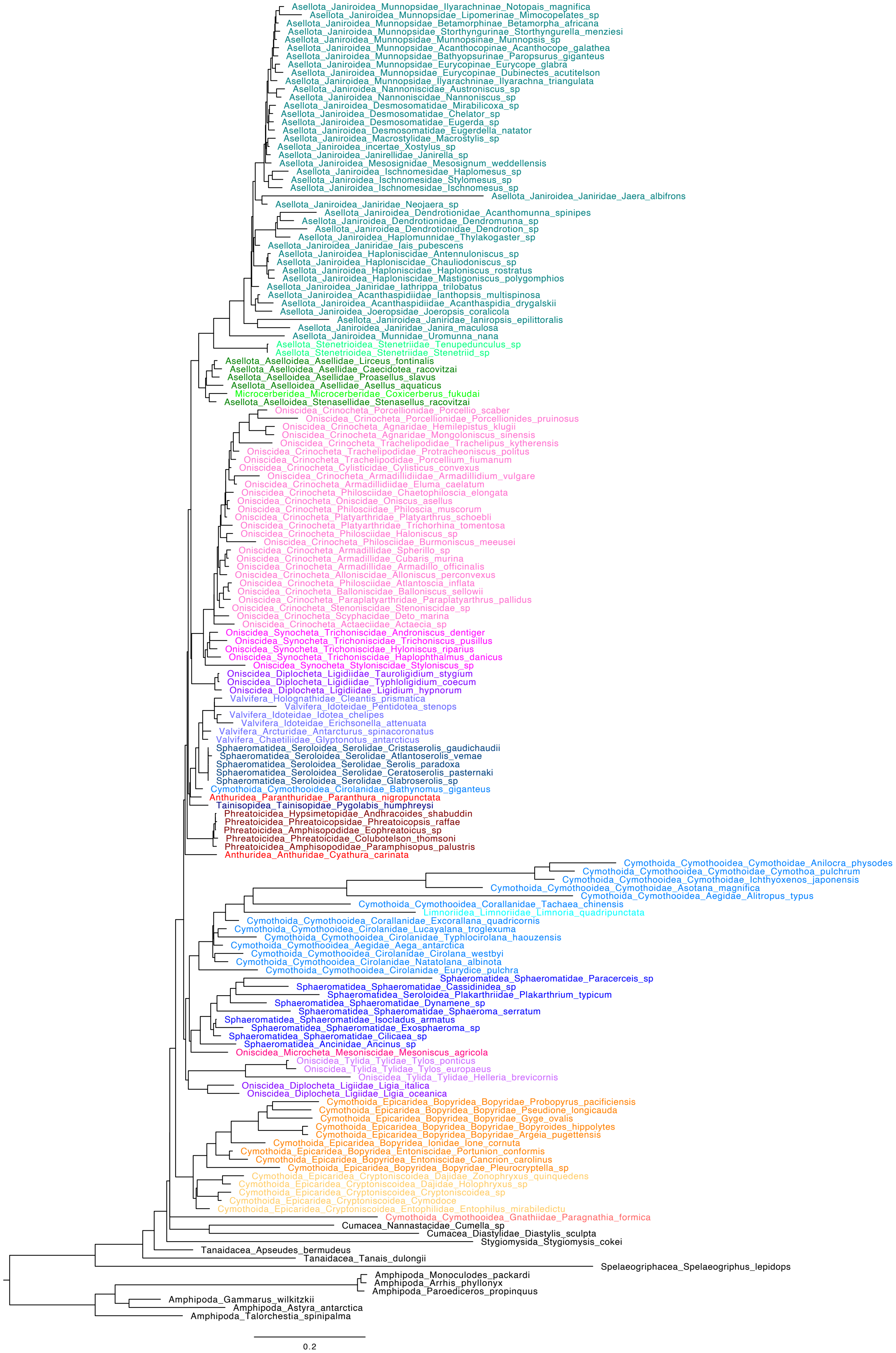

**Supplementary Figure S10:** Phylogeny from ML analysis in *IQ-TREE* of marker-gene dataset 18S, with GHOST model of heterotachy. Branch lengths in substitutions per site.

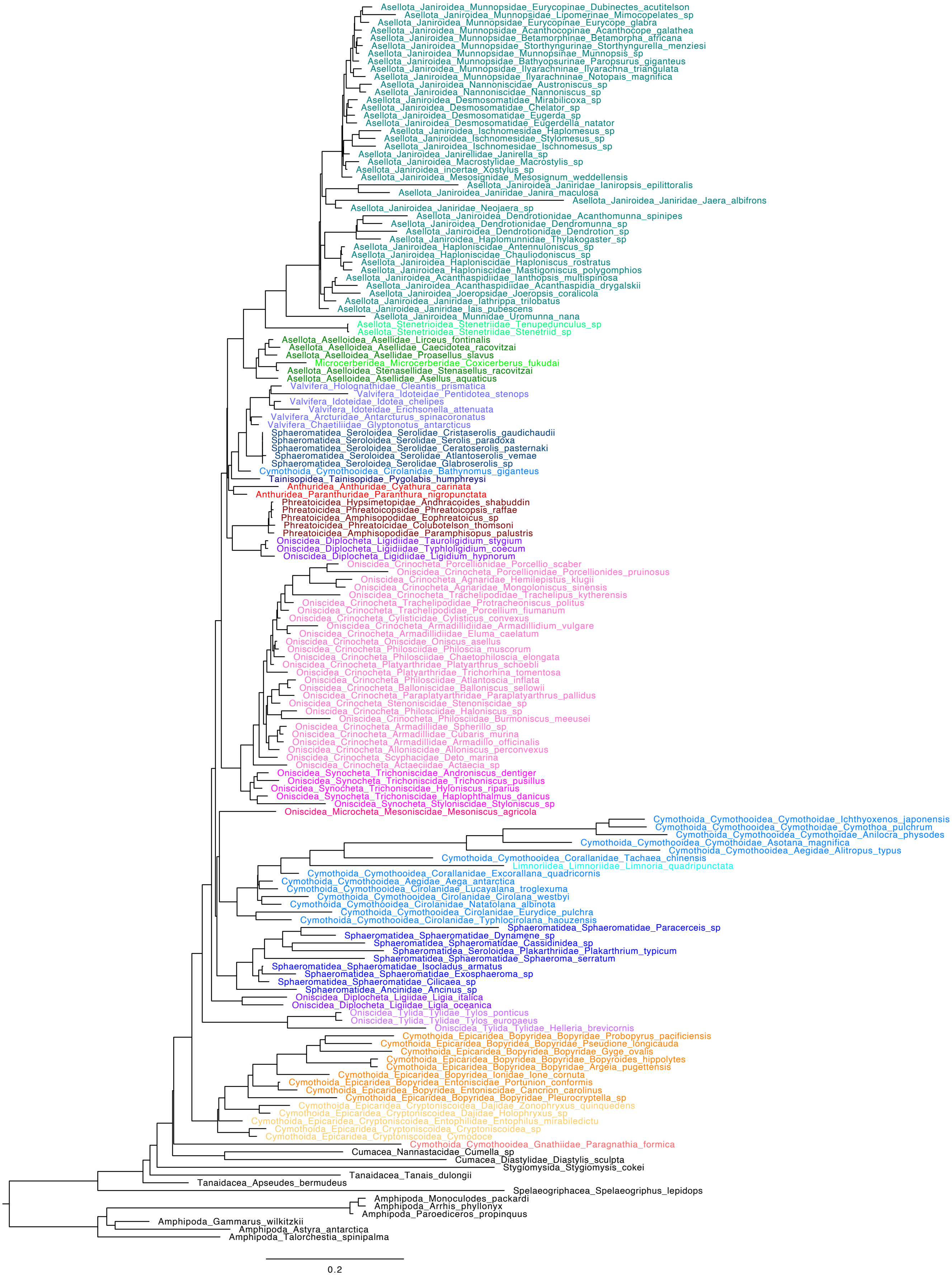

**Supplementary Figure S11:** Phylogeny from ML analysis in *IQ-TREE* of marker-gene dataset 18S, with RY-recoding. Branch lengths in substitutions per site.

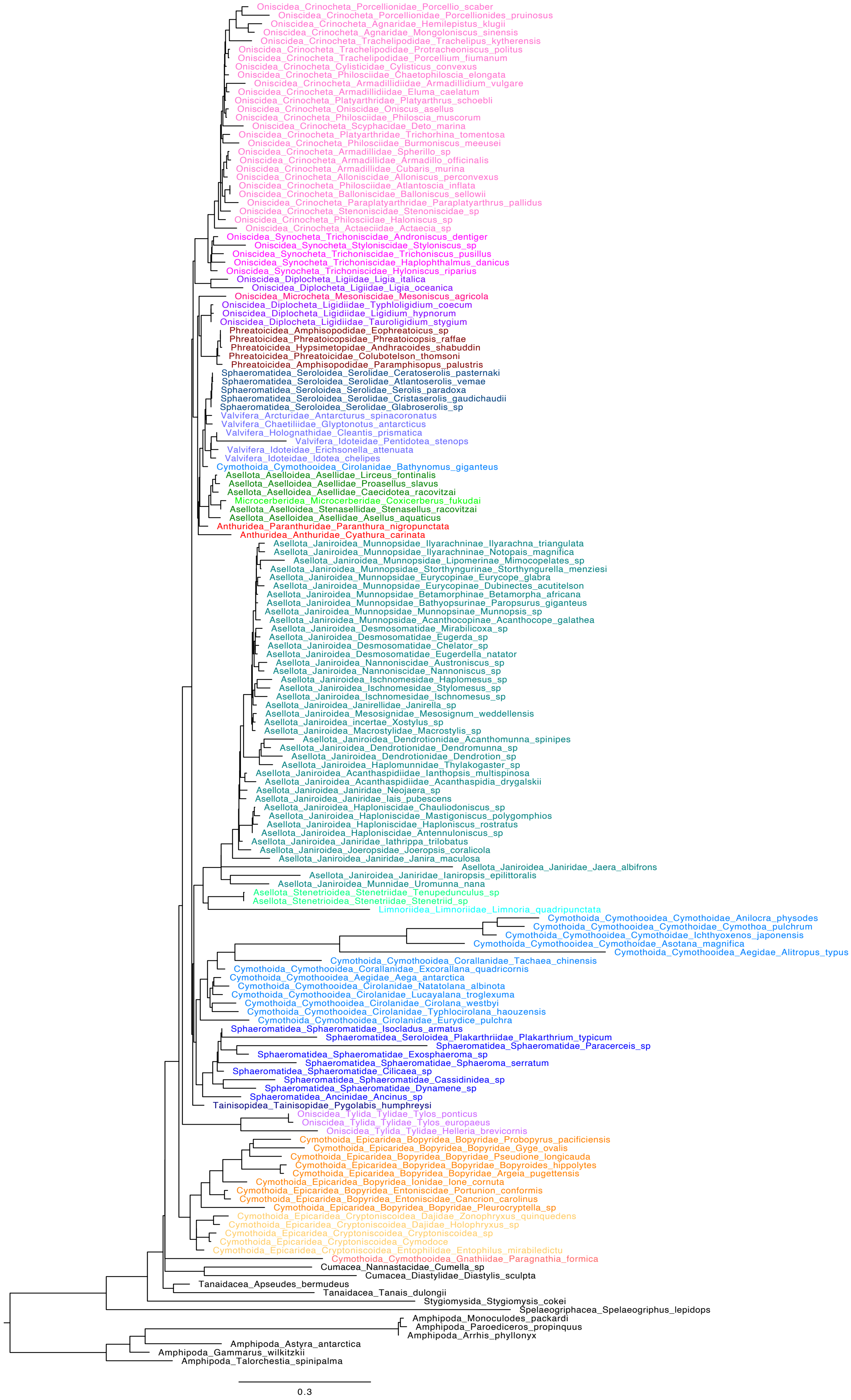

**Supplementary Figure S12:** Phylogenomic dataset, supermatrix of 960 single-copy orthologues, aligned with *FSA* and trimmed, third codon nucleotides positions only, analysed with ML in *IQ-TREE* (a) with GTR+G model of sequence evolution and excluding Synocheta, (b) with chosen best fit model including Synocheta. Branch lengths in substitutions per site.

**(a)**

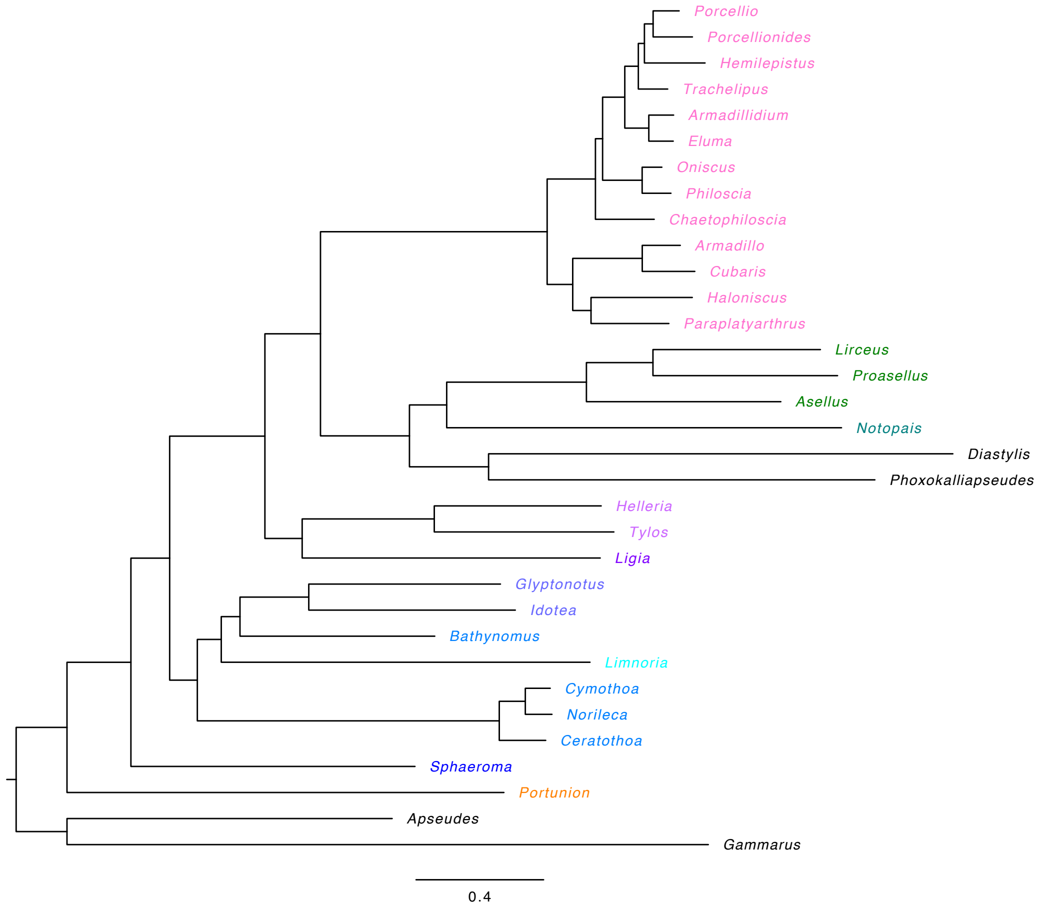


**(b)**

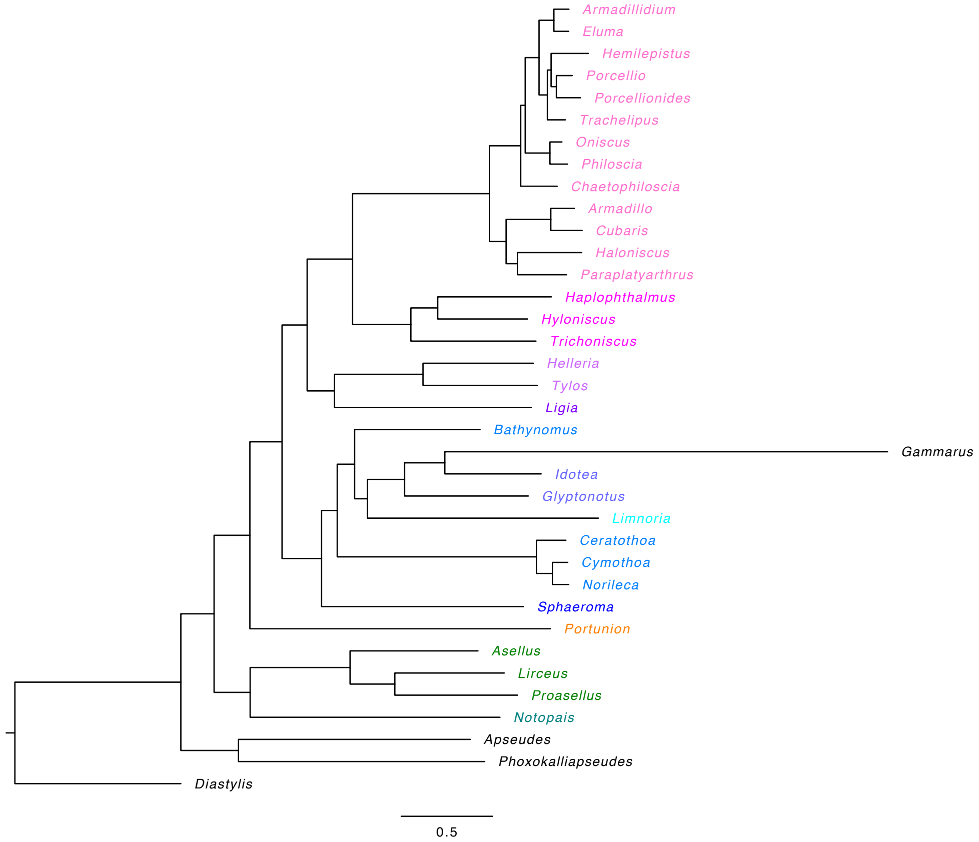

**Supplementary Figure S13:** Dated phylogeny of *BEAST* analysis of marker-gene dataset, with fossil constraints approximating uniform distributions. Branch lengths in millions of years, values at nodes indicate median date estimates in millions of years, node bars indicate 95% highest probability distribution (Bayesian confidence interval measure).

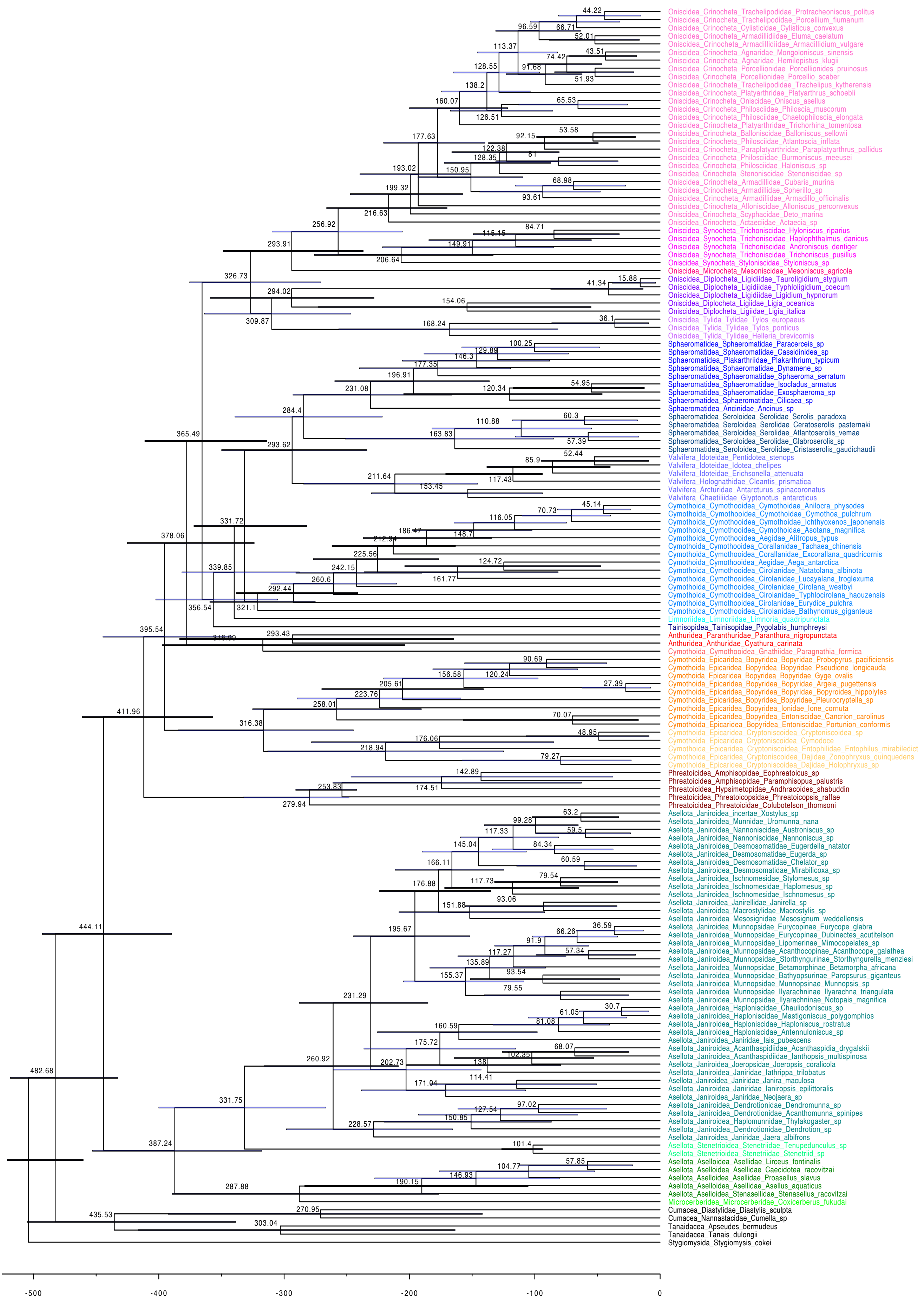
